# The structural organisation of pentraxin-3 and its interactions with heavy chains of inter-α-inhibitor regulate crosslinking of the hyaluronan matrix

**DOI:** 10.1101/2024.08.14.607909

**Authors:** Anokhi Shah, Xiaoli Zhang, Matthew Snee, Michael P. Lockhart-Cairns, Colin W. Levy, Thomas A. Jowitt, Holly L. Birchenough, Louisa Dean, Richard Collins, Rebecca J. Dodd, Abigail R. E. Roberts, Jan J. Enghild, Alberto Mantovani, Juan Fontana, Clair Baldock, Antonio Inforzato, Ralf P. Richter, Anthony J. Day

## Abstract

Pentraxin-3 (PTX3) is an octameric protein, comprised of eight identical protomers, that has diverse functions in reproductive biology, innate immunity and cancer. PTX3 interacts with the large polysaccharide hyaluronan (HA) to which heavy chains (HCs) of the inter-α-inhibitor (IαI) family of proteoglycans are covalently attached, playing a key role in the (non-covalent) crosslinking of HC•HA complexes. These interactions stabilise the cumulus matrix, essential for ovulation and fertilisation in mammals, and are also implicated in the formation of pathogenic matrices in the context of viral lung infections. To better understand the physiological and pathological roles of PTX3 we have analysed how its quaternary structure underpins HA crosslinking via its interactions with HCs. A combination of X-ray crystallography, cryo-electron microscopy (cryo-EM) and AlphaFold predictive modelling revealed that the C-terminal pentraxin domains of the PTX3 octamer are arranged in a central cube, with two long extensions on either side, each formed from four protomers assembled into tetrameric coiled-coil regions, essentially as described by (Noone *et al*., 2022; doi:10.1073/pnas.2208144119). From crystallography and cryo-EM data, we identified a network of inter-protomer salt bridges that facilitate the assembly of the octamer. Small angle X-ray scattering (SAXS) validated our model for the octameric protein, including the analysis of two PTX3 constructs: a tetrameric ‘Half-PTX3’ and a construct missing the 24 N-terminal residues (Δ1-24-PTX3). SAXS determined a length of ∼520 Å for PTX3 and, combined with 3D variability analysis of cryo-EM data, defined the flexibility of the N-terminal extensions. Biophysical analyses revealed that the prototypical heavy chain HC1 does not interact with PTX3 at pH 7.4, consistent with our previous studies showing that, at this pH, PTX3 only associates with HC•HA complexes if they are formed in its presence. However, PTX3 binds to HC1 at acidic pH, and can also be incorporated into pre-formed HC•HA complexes under these conditions. This provides a novel mechanism for the regulation of PTX3-mediated HA crosslinking (e.g., during inflammation), likely mediated by a pH-dependent conformational change in HC1. The PTX3 octamer was found to associate simultaneously with up to eight HC1 molecules and, thus, has the potential to form a major crosslinking node within HC•HA matrices, i.e., where the physical and biochemical properties of resulting matrices could be tuned by the HC/PTX3 composition.

## INTRODUCTION

Hyaluronan (HA) is a large polysaccharide that has a critical role in dictating tissue elasticity, hydration and permeability through its non-covalent interactions with HA-binding proteins (Bano et al., 2018; Richter et al., 2018). HA can also become covalently modified by heavy chains (HCs) of the inter-α-inhibitor (IαI) family of proteoglycans, to form HC•HA complexes (Day and Milner, 2019), a process mediated by the protein TSG-6 (Rugg et al., 2005; Sanggaard et al., 2008).

Pentraxin-3 (PTX3), an octameric protein (Inforzato et al., 2010, 2008) with diverse biological roles (Garlanda et al., 2018), has been found to play an important structural role via its crosslinking of HC•HA complexes (Baranova et al., 2014), e.g., in ovulation (Camaioni et al., 2018). In this context, HC•HA and PTX3 are both required for stabilisation and expansion of the cumulus matrix that forms around the oocyte and is critical for fertilisation in vivo (Briggs et al., 2015; Salustri et al., 2004; Sato et al., 2001; Varani et al., 2002; Zhuo et al., 2001). The cumulus matrix has unusual material properties, with the cumulus-oocyte complex (COC) being the softest elastic tissue so far described (Chen et al., 2016). However, the molecular basis of these properties is not understood. In addition, HC•HA/PTX3 complexes extracted from human amniotic membrane (He et al., 2009; Zhang et al., 2012) have potent anti-inflammatory and wound healing activities, through their promotion of cell-cell interactions and signalling (Tseng, 2016; Zhu et al., 2020). HC•HA complexes also form during disease processes, including in the lung during asthma (Forteza et al., 2007; Lauer et al., 2015), ozone-induced airway hyperresponsiveness (Stober et al., 2017), and following infection with influenza A (Bell et al., 2019; Tang et al., 2024) and SARS-CoV-2 (Queisser et al., 2021); with upregulation of PTX3 expression reported in some of these cases (Brunetta et al., 2020; Koussih et al., 2021).

There are up to 5 different HCs (HC1, HC2, HC3, HC5 and HC6; also known as ITIH1, 2, 3, 5 and 6) that can become covalently associated with HA, and there is evidence that the composition of HC•HA complexes is context and tissue specific (Day and Milner, 2019). The crystal structure of HC1 (where the mature protein corresponds to residues 35-672 of the pre- protein; see UniProt P19827) was recently determined, defining the canonical fold of the ITIH family (Briggs et al., 2020); this comprises a large 16-stranded β-sandwich domain composed of sequences from the N- and C-terminal portions of the protein (HC-Hybrid2), along with a region that resembles an integrin β-subunit because it contains a von Willebrand Factor A (vWFA) domain (including a Metal-Ion Dependant Adhesion Site (MIDAS)) and an associated hybrid domain (HC-Hybrid1). HCs attach covalently to HA via a C-terminal Asp residue (Zhao et al., 1995), however, the last 20 amino acids of HC1 were not visible in the electron density and therefore were presumed to be unstructured or conformationally dynamic (Briggs et al., 2020). HC1 was found to form a metal-ion-, and MIDAS-, dependant dimer, as shown by analytical ultracentrifugation (AUC) and small-angle X-ray scattering (SAXS) techniques. While these data (Briggs et al., 2020) provide direct evidence for HC-HC interactions, which had been suspected based on electron microscopy (EM) of HC•HA purified from the synovial fluids of patients with rheumatoid arthritis (Day and Milner, 2019; Yingsung et al., 2003), the HC1-HC1 interaction is weak (i.e., *K*_D_ ≍ 40 µM) (Briggs et al., 2020), and therefore unlikely to act as a major crosslinking node between HC-modified HA chains.

PTX3 is made up of eight identical protomers, each comprised of a 161 amino acid N-terminal domain (residues 18-178 in Uniprot P26022) and a 203 amino acid C-terminal ‘pentraxin’ domain (residues 179-381), connected via a disulphide bond network (Inforzato et al., 2010, 2008). The N-terminal domain (N_PTX3) is responsible for the covalent oligomerisation of PTX3 into tetramers through inter-chain disulphide bridges via three cysteine residues (47, 49 and 103); the N-terminal end of N_PTX3 is thought to be intrinsically disordered (Doni et al., 2019; Inforzato et al., 2010; Noone et al., 2022). The C-terminal domain of PTX3 (C_PTX3) is homologous to the short pentraxins C-reactive protein (CRP) and serum amyloid P component (SAP) (Inforzato et al., 2010; Scarchilli et al., 2007), and contains a single N-linked glycan at Asn-220 (Inforzato et al., 2006). Inter-chain disulphide bonds between Cys318 and Cys319 link together the PTX3 tetramers to form octamers (Inforzato et al., 2010, 2008), however, these vicinal cysteine residues can alternatively participate in intra-chain disulphides in some of the protomers (Inforzato et al., 2008).

As described above, PTX3 contributes to the formation of the cumulus matrix prior to ovulation (Salustri et al., 2004; Varani et al., 2002). In the absence of PTX3, while HC•HA production is not impaired, the HC•HA complexes fail to incorporate into (and crosslink) the matrix, revealing a critical role for PTX3 in COC expansion (Salustri et al., 2004). This crosslinking activity of PTX3 is reliant on its oligomerisation state, such that the tetrameric N_PTX3 domain, and mutant forms of the full-length protein that are tetramers, but not those that are dimers, are able to rescue the expansion of COCs from PTX3-deficient mice (Inforzato et al., 2008; Scarchilli et al., 2007). The rescue of COC matrix cross-linking by N_PTX3 and the interaction of this construct with IαI led to the conclusion that the binding sites for HCs are localised within the N-terminal domain of PTX3 (Inforzato et al., 2010; Scarchilli et al., 2007), however, no interaction experiments with HCs have been performed and the stoichiometry is not known. Importantly, the interaction of PTX3 with HC•HA is tightly regulated (Baranova et al., 2014). In this regard, PTX3 does not interact with preformed HC•HA complexes at pH 7.4 and only becomes incorporated into the HA network if PTX3 interacts with IαI prior to the formation of HC•HA.

Previously, a low-resolution solution structure for PTX3 was determined from SAXS, leading to the generation of a molecular model in which the 8 C-terminal pentraxin domains were proposed to assemble into a central body (in a pseudo cubic arrangement) with regions comprised of 4 N_PTX3 domains (arranged asymmetrically as tetrameric coiled-coils) situated to either side (Inforzato et al., 2010); this model, was consistent with the disulphide bond network determined for PTX3 (Inforzato et al., 2008). More recently, a hybrid model of PTX3 has been generated (Noone et al., 2022), providing a high-resolution cryo-EM structure (2.5 Å) for the central ‘cube’ (with the eight pentraxin domains having D4 symmetry) in combination with AlphaFold modelling of the N-terminal domains, also predicting that they form tetramers involving coiled-coli regions, with flexible hinges (7ZL1.pdb); however, in Noone *et al*. (Noone et al., 2022), the N_PTX3 domains were much more elongated, and with a symmetric arrangement, compared to the earlier model (Inforzato et al., 2010).

Here, we have determined a structure for PTX3 based on X-ray crystallography, cryo-EM and molecular modelling. This has confirmed the overall structural organisation described for PTX3 by Noone and colleagues (Noone et al., 2022). Our EM-density additionally allows for the modelling of a region of the N-terminal domain, demonstrating that it is a helical tetrameric coiled-coil. Our PTX3 model has been validated by SAXS analysis on a range of constructs (full-length and truncated). In addition to structural studies, we have also extensively investigated the interaction of PTX3 with HC1 (as an archetypal heavy chain) by biophysical methods, making use of a ‘half-PTX3’ construct with only one N-terminal extension. While there is no interaction between HC1 and PTX3 at pH 7.4 (consistent with previous findings for HC•HA (Baranova et al., 2014)), the proteins did interact strongly at pH 5.5 (with a low nM *K*_D_ value), providing a mechanism for pH-mediated regulation. Moreover, we show that the PTX3 octamer has the potential to bind eight HC1 molecules simultaneously and propose a novel interaction site based on differential binding of HC1 to two polymorphic variants of PTX3. Overall, this study provides new molecular insights into a major crosslinking node (HC•HA/PTX3) present within HA matrices formed during health and disease, based on the regulated crosslinking of HC•HA complexes via multimeric PTX3-HC interactions.

## RESULTS

### Structural analysis of PTX3

We initially determined a high-resolution X-ray crystal structure of an individual C-terminal (pentraxin) domain of human PTX3 (Figure 1), denoted here as C_PTX3 (see definitions in Figure S1). The structure, comprising residues 178-381 in the preprotein, was expressed and purified as previously reported (Inforzato et al., 2010), and crystallised, yielding an X-ray structure to 2.4 Å resolution (Table 1; 8PVQ.pdb). The C_PTX3 monomer (Figure 1A) adopts a fold with a β-sandwich architecture, which is typical of other pentraxin domains (Emsley et al., 1994; Suzuki et al., 2020). Residues 316-320, which form part of the loop containing the vicinal cysteines (C317 and C318) required for oligomerisation, were modelled as these were unresolved in the electron density, likely due to flexibility (Figure S2). This is consistent with data that indicates that there is heterogeneity in the disulphide bond formation within this region (Inforzato et al., 2008), where both inter- and intra-protomer disulphide bonds can form. Superposition of our C_PTX3 structure with a monomer of the C-terminal domain from the model proposed by Noone *et al*. (Noone et al., 2022) revealed a backbone root mean square deviation (RMSD) of 0.9 Å.

**Figure 1.**
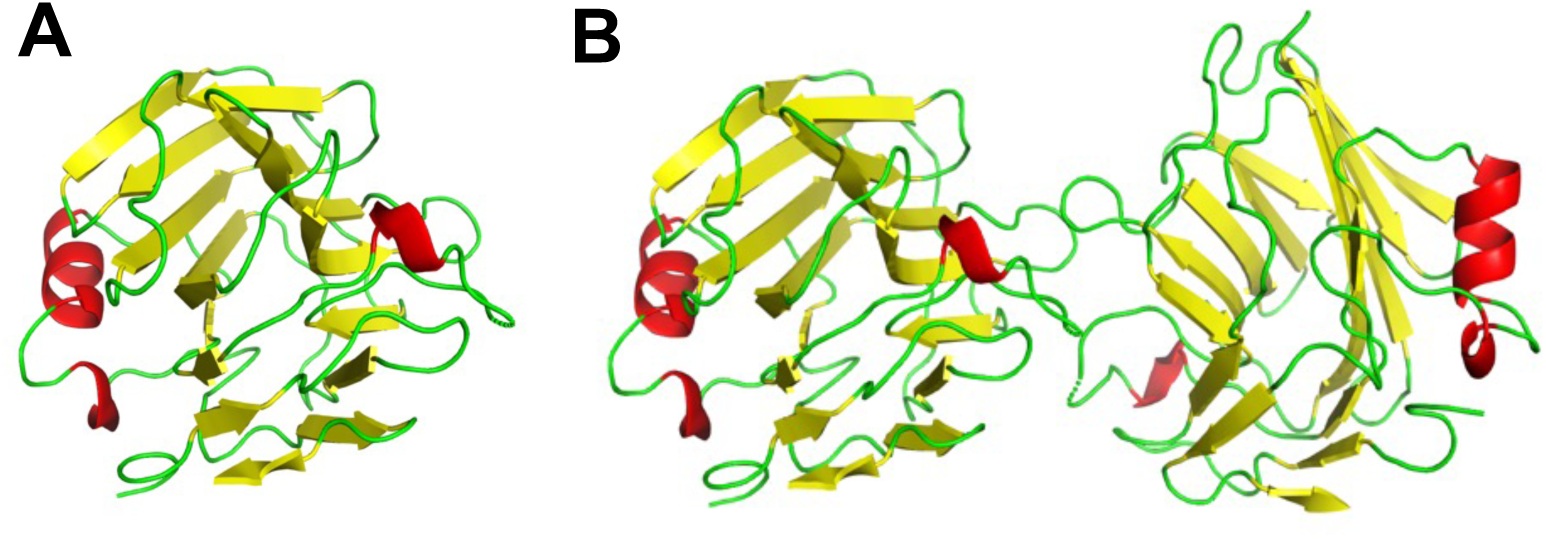
Structure of the pentraxin domain of PTX3. (**A**) X-ray crystal structure (8PVQ) of the C_PTX3 monomer resolved to 2.4 Å with α-helices and β-sheets are shown in red and yellow, respectively. (**B**) Two adjacent C_PTX3 monomers from the crystallographic unit cell; these pentraxin domains are present in a relative orientation that is essentially identical to that seen for adjacent pentraxin domains in the cryo-EM map (Figure 3A).

**Table 1.** Data collection and refinement statistics for C_PTX3 crystal structure. Statistics for the highest-resolution shell are shown in parentheses.

|  | **C_PTX3 (8PVQ.pdb)** |
| --- | --- |
| Wavelength | 0.98 Å |
| Resolution range | 45.33-2.43 (2.52-2.43) |
| Space group | H 3 2 |
| Unit cell | 138.0, 138.0 69.6, 90, 90, 120 |
| Total reflections | 62,653 (5,655) |
| Unique reflections | 9,663 (950) |
| Multiplicity | 6.5 (6.0) |
| Completeness (%) | 99.54 (99.37) |
| Mean I/sigma(I) | 8.66 (1.08) |
| Wilson B-factor | 62.74 |
| R-merge | 0.1040 (1.504) |
| R-meas | 0.1131 (1.647) |
| R-pim | 0.04392 (0.6641) |
| CC1/2 | 0.998 (0.661) |
| CC* | 0.999 (0.892) |
| Reflections used in refinement | 9,637 (944) |
| Reflections used for R-free | 483 (53) |
| R-work | 0.2165 (0.3624) |
| R-free | 0.2633 (0.3763) |
| CC (work) | 0.952 (0.757) |
| CC (free) | 0.877 (0.718) |
| Number of non-hydrogen atoms | 1567 |
| macromolecules | 1546 |
| ligands | 0 |
| solvent | 21 |
| Protein residues | 200 |
| RMS (bonds) | 0.009 |
| RMS (angles) | 0.80 |
| Ramachandran favoured (%) | 96.43 |
| Ramachandran allowed (%) | 3.06 |
| Ramachandran outliers (%) | 0.51 |
| Rotamer outliers (%) | 1.82 |
| Clashscore | 4.26 |
| Average B-factor | 69.94 |
| macromolecules | 69.97 |
| solvent | 67.71 |

To expand our structural understanding of PTX3 beyond the C_PTX3 monomer, cryo-EM data (Dataset 1; see Methods) were generated for recombinant full-length PTX3 (Figure 2), showing a cube-like arrangement of eight protomers with D4 symmetry forming the Octameric C-terminal Domain (OCD_PTX3); see Figure S3 for SDS-PAGE analysis of PTX3 proteins used in the study. The RMSD of a monomeric unit within the OCD_PTX3 cryo-EM structure compared with the monomeric PTX3 X-ray structure is 0.704 Å (Figure S4). This allowed us to dock the crystal structure for C_PTX3 into the cryo-EM density followed by manual rebuilding and refinement, as described in the Methods section, i.e., generating a ‘model’ for OCD_PTX3 that was informed by the crystal structure. The cryo-EM density also resolved 26 residues (per protomer) of the N-terminal domain of PTX3 (Figure 2C-E) as compared to 19 residues from a low-pass filtered map to ∼5 Å solved for PTX3 previously (7ZL1.pdb (Noone et al., 2022)). Residues 150-177 of the Tetrameric N-terminal Domain of PTX3 (TND_PTX3) that are resolved in our cryo-EM structure are highlighted in Figure 2D-E showing how four protomers come together to form a tetrameric coiled-coil arrangement. The overall resolution of our cryo-EM structure is 3.3 Å, with higher resolution (∼2.5 Å) within the OCD, comprised of eight pentraxin domains, and lower resolution in the N-terminal region (∼4 Å); Figure S5 and Figure S6, respectively, show the corresponding local resolution map and its validation.

**Figure 2.**
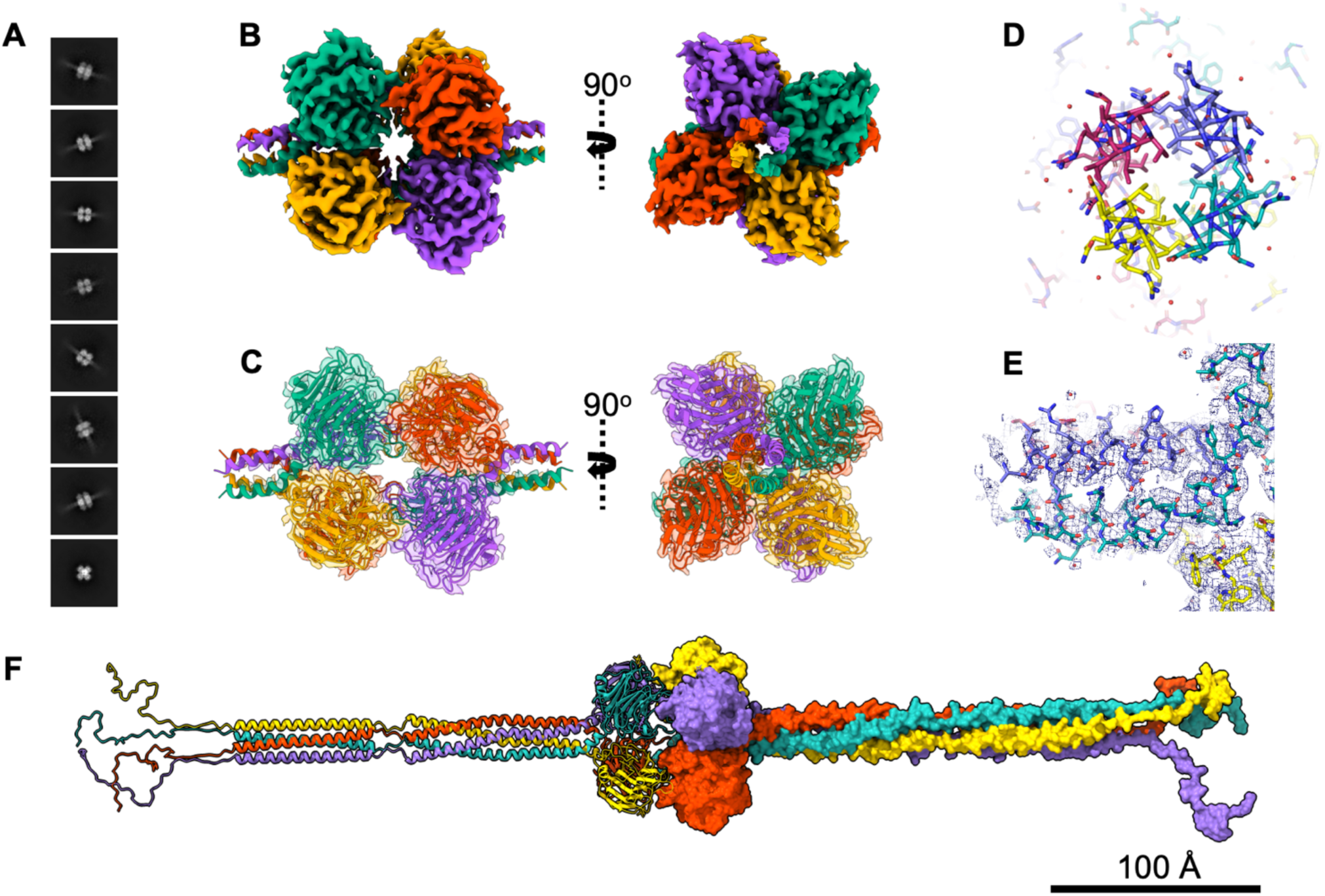
Cryo-EM structure of PTX3. (**A**) Selected 2D classes representing ‘side-views’ of PTX3 particles, in which the N-terminal tetrameric domains (TND_PTX3) are apparent as diffuse extensions emanating from the central OCD_ PTX3; boxes are 38 nm × 38 nm in size. (**B**) Cryo-EM map of OCD_PTX3 (with D4 symmetry) including a region of the interface with TND_PTX3 (26 residues), resolved to 3.3 Å. The four colours (turquoise, purple, red and yellow) represent the different protomers within each tetramer of PTX3. (**C**) Tertiary structure of residues 150-381 built into the EM map, of the full OCD_PTX3 plus 26 adjacent residues of the TND_PTX3. (**D**, **E**) Top and side views, respectively, of the structurally resolved coiled-coil region at the interface of the C- and N-terminal domains. (**F**) Model of PTX3, where the OCD_PTX3 (residues 178-381) and residues 150-177 of the TND_PTX3 were determined here (in **C**) and residues 18-149 were modelled by AlphaFold. Residues 18-46 were relaxed using the ‘Model Loops’ tool in Chimera; the left-hand tetramer of PTX3 is shown as a ribbon representation and the right-hand tetramer as a surface, both coloured as in **B**.

The remaining tetrameric N-terminal domain, which was not sufficiently resolved by EM density, was modelled using AlphaFold (Jumper et al., 2021), being built onto our high-resolution cryo-EM structure (see Methods). The final model generated is shown in Figure 2F, where this has an extended disordered region at its N-terminal end (residues 18-46), followed by a short ‘linear’ section (residues 47-54) containing the cysteine residues (C47 and C49) that form inter-chain disulphide bonds, and two regions of tetrameric coiled-coil (residues 55-100 and 107-172), separated by a short sequence containing C103, which also forms an inter-protomer disulphide.

Given that our overall PTX3 structure/model is very similar to that of Noone *et al*. (Noone et al., 2022), here we have focused on insights not covered previously and how we validated our model experimentally. The high resolution of the cryo-EM structure within the central OCD_PTX3 region enabled us to resolve salt bridges formed between protomers and identify that these interactions have a key role in promoting the stabilisation of the oligomeric structure of PTX3 (Figure 3). Tetramers are stabilised by interactions between E180 and K214 in adjacent chains such that salt bridges are formed at each of the interfaces between the four protomers (Figure 3B); within the same protomer, E180 and K214 are located on oppositive faces of the pentraxin domain. There may also be an additional salt bridge that stabilises the tetramer, where this links the carboxylate of S381 to the sidechain of R172 (i.e., the last residue of the second coiled coil region in TND) in an adjacent protomer. However, the electron density is not well resolved for this C-terminal residue.

**Figure 3.**
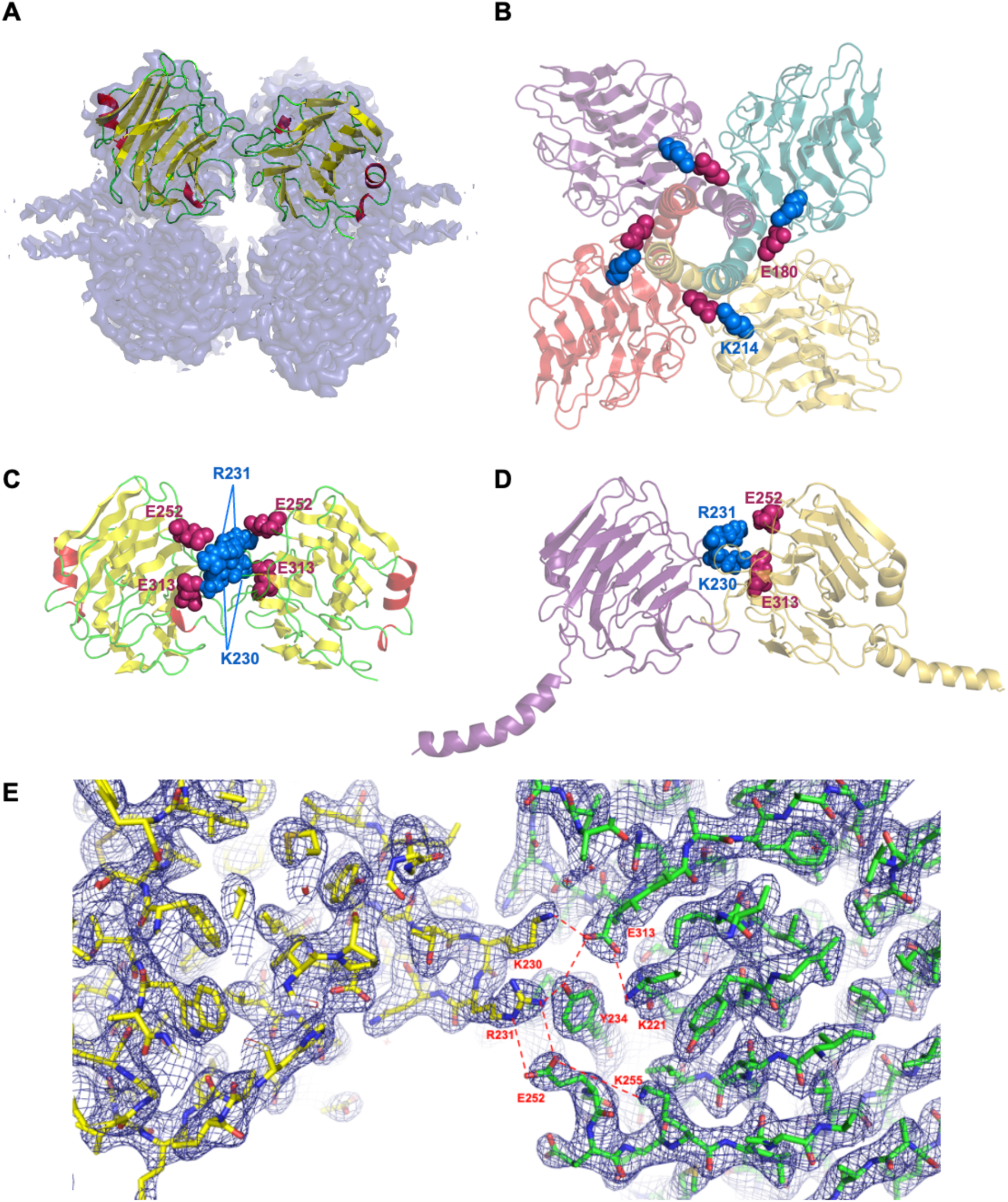
Salt bridges promote the formation of the octameric structure of PTX3. (**A**) Two adjacent C_PTX3 monomers from the crystallographic unit cell (α-helices in red and β-strands in yellow) superimposed on the cryo-EM map for PTX3 (grey). (**B**) The tertiary structure for PTX3 (determined here) showing 4 protomers (coloured: magenta, blue, yellow and orange) where four salt bridges between E180 and K214 residues stabilise the formation of the tetramer; Glu and Lys residues in all 4 protomers are shown in space-filling representation. (**C**) Adjacent C_PTX3 monomers from **A** (rotated ∼90° forwards around the x-axis) showing the residues that form additional salt bridges between the tetramers (to stabilise the octamer). For each pair of protomers, four salt bridges are formed, i.e., from K230, R231, E252 and E313 in one protomer to E313, E252, R231 and K230, respectively, in the other. (**D**) A pair of protomers from the PTX3 structure with the C_PTX3 domains in the same orientation as in **C**. For clarity, salt bridges are only shown between K230 and R231 (in the magenta protomer) and E313 and E252 (in the yellow protomer), respectively. (**E**) Electron density from the crystal structure of C_PTX3 showing the network of interactions (red) formed between two monomers; essentially identical interactions are present in the PTX3 cryo-EM structure, i.e., linking together two tetramers into an octamer.

Importantly, residues K230 and R231 in one protomer form ionic interactions with E313 and E252, respectively, in an adjacent protomer, linking together the two tetramers to stabilise the octamer (Figure 3C,D). The reciprocal pair of salt bridges (E252-R231 and E313-K230) are also formed, meaning that there are four salt bridges between each pair of protomers. These salt bridges are present in both the cryo-EM (Figure 3D) and crystal structures (Figure 3C), where in the latter two adjacent C_PTX3 monomers in the unit cell (Figure 1B) adopt an orientation that closely matches that of adjacent pentraxin domains in the cryo-EM map (Figure 3A). The conformation of E313 is stabilised through intra-protomer ionic interaction with K221 and a hydrogen bond to Y234 (as can be seen from the electron density for the crystal structure; Figure 3E). It is possible that E252 may also be stabilised by K255 in a similar manner, albeit with alternating salt brides between R231-E252 and E252-K255 being formed.

The presence of these various salt bridges within the crystal lattice for the isolated C_PTX3 domain (Figure 3C,E), as well as in PTX3 in its native octameric state (Figure 3D), indicates their importance in the formation of the octamer. Here it can be envisaged that the E180-K230 and K214-R231 interactions generate sixteen salt bridges driving the association of the two tetramers ahead of disulphide bond formation between C317 and C318.

### Validation of the PTX3 model by Small-Angle X-ray Scattering (SAXS)

PTX3 has a single N-linked glycosylation site (N220) located on the C-terminal pentraxin domain of each protomer. Glycan analysis of the PTX3 construct expressed in Chinese hamster ovary (CHO) cells (and used for the cryo-EM work), identified the glycan of highest abundance (Figure S7), which was modelled onto our structure of PTX3 (Figure 4Ai). To experimentally validate the model of glycosylated PTX3, SEC-SAXS data (in solution) were collected for the PTX3 protein expressed in CHO cells (Figure 4Ai; Figure S8; Table S1). The normalised Kratky plot (Rambo and Tainer, 2011) has a shape consistent with that of an elongated multidomain protein (Figure S8B), composed of a well-folded domain and a flexible extension, with a maximum dimension (*d*_max_) of at least 430 Å (Figure S8D,E; Table S1). When comparing the experimental SAXS data for PTX3 with the theoretical scattering of our glycosylated model, the curves superimpose well (Figure 4Ai) indicating that the model is an accurate representation of PTX3. Although visually the fit is good, the normalised χ^2^ is high (χ^2^ = 38.8), due to poorer fit quality in the mid-q region. This likely reflects a contribution from the flexibility of the TND_PTX3 (see below), as well as the presence of complex glycosylation states; only the most common out of five abundant glycans (Figure S7) was modelled, in the single most favourable rotamer.

**Figure 4.**
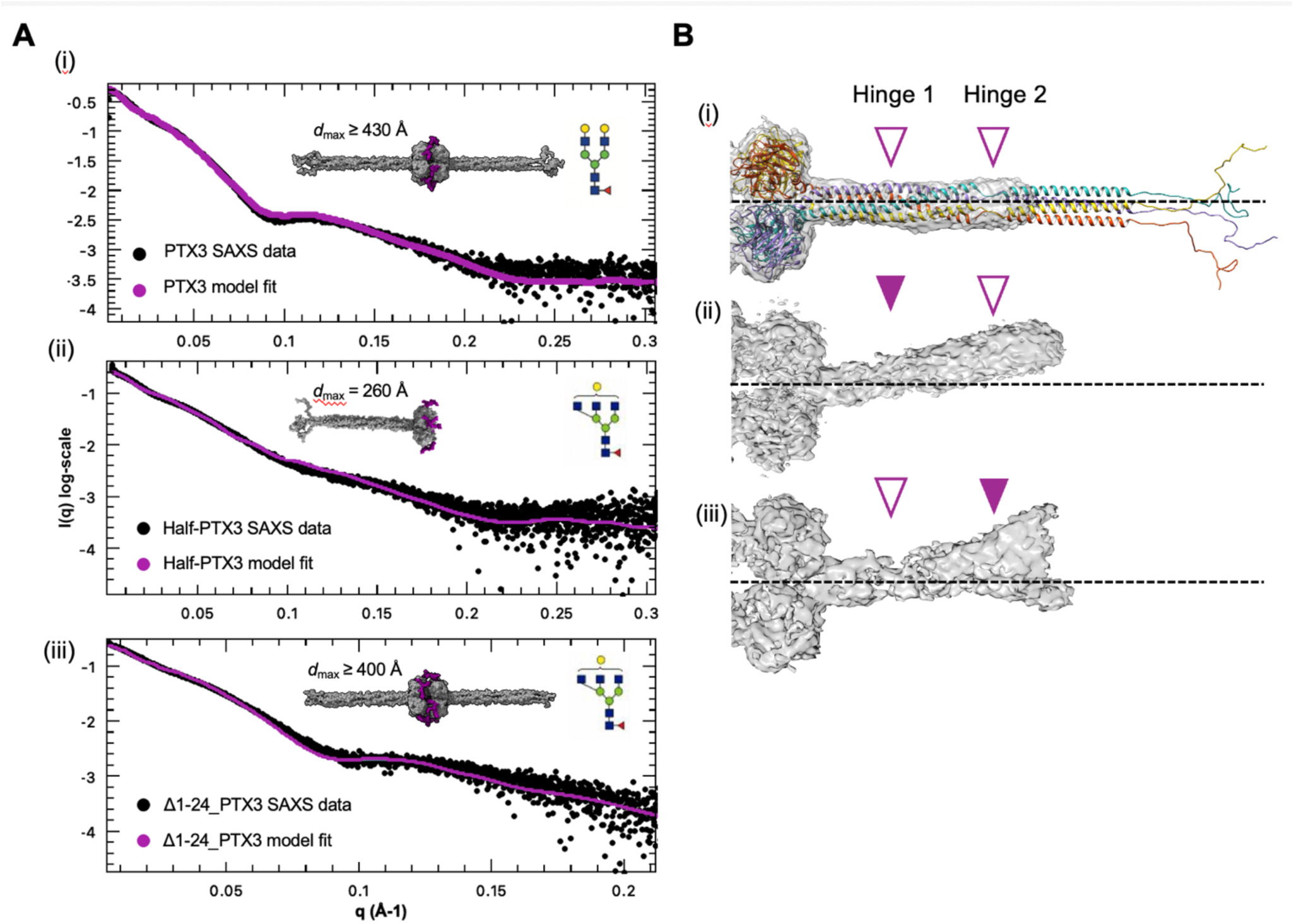
SAXS of PTX3 and 3D variability analysis from cryo-EM revealing flexibility of the tetrameric N-terminal domain. (**A**) SEC-SAXS analysis of PTX3 constructs. Black dots: SAXS data; purple lines: theoretical scattering curves for the respective models in **i**-**iii** with most abundant glycan (schematic; see Figure S7 for details) modelled in purple: (**i**) PTX3 expressed in CHO cells; (**ii**) Half-PTX3 expressed in HEK Expi293F cells; (**iii**) Δ1-24_PTX3 expressed in HEK Expi293F cells. (**B**) 3D variability analysis (3DVA) of the low resolution cryo-EM maps for a tetramer of PTX3 with arrowheads showing points of flexibility. (**i**) Cryo-EM map, showing one tetramer, where the N-terminal region is unflexed (grey surface representation) overlayed with the PTX3 model (coloured ribbon representation). (**ii**-**iii**) Maps of the N-terminal domains with flexed structures, with purple arrowheads showing the positions of ‘hinges’ where bending occurs: (**ii**) map in which only Hinge 1 is flexing (solid arrow); (**iii**) map where only Hinge 2 is flexing (solid arrow); here density can be seen in two different positions reflecting molecules within the dataset with bend in opposite directions.

Two truncated constructs of PTX3 were made (see Figure S1) and investigated by SEC-SAXS. In the first, cysteines C317 and C318 (that form interchain disulphides linking the two tetramers of PTX3 together (Inforzato et al., 2010, 2008)) were mutated to serines to generate a ‘Half-PTX3’ construct (i.e., a tetramer composed of 4 full-length protomers). In the second, the first 24 amino acids of the mature protein (corresponding to residues 18-41 in the preprotein) were deleted, which are predicted to be disordered based on our modelling (Figure 2C), and this was denoted Δ1-24_PTX3. These constructs were expressed in HEK Expi293F cells and, unsurprisingly, were found to have a different glycosylation pattern to the CHO expressed PTX3 (Figure S7). Models for Half-PTX3 and Δ1-24_PTX3 were made based on our PTX3 structure, where the highest abundance glycan (from PTX3 expressed in HEK Expi293F cells) was modelled onto the C_PTX3 domains, and SEC-SAXS data were collected for the corresponding constructs (Figures 4Aii and 4Aiii, respectively). The experimental data fit reasonably well to the glycosylated models (Figure 4A), but again with large normalised χ^2^ values (χ^2^ of 37.5 for Half-PTX3 and 21.4 for Δ1-24_PTX3) likely due to the TND_PTX3 flexibility and/or the presence of complex glycosylation states. The experimental *d*_max_ of 260 Å for the Half-PTX3 construct (which is the most reliable value of the 3 constructs given the detector distance used in SAXS data collection) fits well with our Half-PTX3 model. Therefore, the full-length PTX3 would be expected to have a maximum length of ∼520 Å, which is somewhat longer than the model in Noone *et al*. (Noone et al., 2022) where the N-terminal peptides (residues 18-46), while disordered, fold back along the first coiled coil. Thus, the extended orientation for this region in our model seems a more likely arrangement and is consistent with our SAXS analysis.

### The coiled-coil region of the Tetrameric N-terminal Domain of PTX3 has flexible hinges

To characterise PTX3 in solution, we analysed the SAXS data for the PTX3, Half-PTX3 and Δ1-24_PTX3 constructs (Figure S8). The normalised Kratky plot (Figure S8B) shows two distinct peaks for the full-length PTX3 and Δ1-24_PTX3, consistent with two discrete folded regions where the initial peak corresponds to the non-flexible OCD region and the second smaller peak to more flexible TND regions. The Half-PTX3 has a less prominent second peak, consistent with it having only one TND.

Flexible proteins are expected to exhibit a plateau the in either the q^2^ (Kratky-Debye) or q^3^ (SIBYLS) plot and less flexible proteins are expected to exhibit a plateau in the q^4^ (Porod-Debye) plot (Rambo and Tainer, 2011). Given that Half-PTX3 is simpler in structure, compared to both Δ1-24_PTX3 and the full-length protein, it is more straightforward to interpret in such a flexibility analysis. In this regard, Figure 8F shows a plateau for Half-PTX3 in the q^3^ before the q^4^ plot (left- and right-hand panels, respectively), indicating that this construct is folded but flexible in solution. Moreover, the Δ1-24_PTX3 plateaued before PTX3 in the q4 plot (data not shown), indicating it is less flexible than the full-length protein consistent with the N-terminal 24 amino acids being unstructured and able to adopt a range of (likely extended) conformations.

To further characterise PTX3 flexibility, we implemented a 3D variability analysis (3DVA) on a different cryo-EM dataset (Dataset 2; Table S2) as a final validation of our hybrid model. Using 14,376 particles representing only side views, we were able to obtain low resolution (<8 Å) density for ∼50% of the length of the TND_PTX3 coiled-coil region (residues 86-172). 3DVA on this dataset (Figure 4B) demonstrate there are two points of major flexibility in the TND of PTX3 (Hinge 1 and Hinge 2; denoted by the purple arrows in Figure 4B) as well as flexibility close to the interface of the OCD and TND. Hinge 1 is centred around P142, which has previously been described by Noone *et al*. as a stutter in the heptad repeat (Noone et al., 2022). Hinge 2 is located where AlphaFold predicts a ‘linear’ unstructured region that interconnects the two coiled-coil regions (residues 101-106; including C103 that forms an inter-protomer disulphide bond. Moreover, the position of the Hinge 2 region largely coincides with the point beyond which the electron density is no longer resolved in the < 8Å cryo-EM map (Figure 4B). This indicates that Hinge 2 may be a region of particular flexibility. Overall, the SAXS data and 3DVA together provide good validation for our PTX3 structure, as well as novel insights into the conformational dynamics of this highly elongated octameric protein.

### HC1 binds to PTX3 under acidic conditions and undergoes a pH-dependent conformational change

Previous work showed that PTX3 does not interact with pre-formed HC•HA complexes (immobilised on sensor chip surfaces) at pH 7.4 (Baranova et al., 2014). Consistent with this we have found that there is no association between PTX3 and HC1 (as an archetypal HC) in solution at this pH (as assessed by size exclusion chromatography (SEC); data not shown). However, PTX3 has been found to bind to some ligands, e.g., fibrin/fibrinogen and plasminogen, with a higher affinity at acidic pH (Doni et al., 2015). Therefore, we investigated the effect of pH on the interaction of PTX3 with HC1.

To facilitate interaction analysis, we designed a construct in which amino acids 316-322 (forming a flexible loop in the crystal structure of C_PTX3; Figure S2) were replaced by a sequence of 7 histidine residues. As in Half-PTX3, this removes C317 and C318 (which form interchain disulphides linking together two tetramers in PTX3). The construct, denoted Half-PTX3-H_7_ (Figure S1), can be attached (via its four H_7_ loops) to supported lipid bilayers (SLBs) made from DOPC lipids and a small fraction of lipid analogues presenting (Ni^2+^-NTA)_3_ moieties (Figure 5A); this provides for a well-defined orientation, with tetrameric N-terminal domains pointing into the solution phase and available for ligand binding.

**Figure 5.**
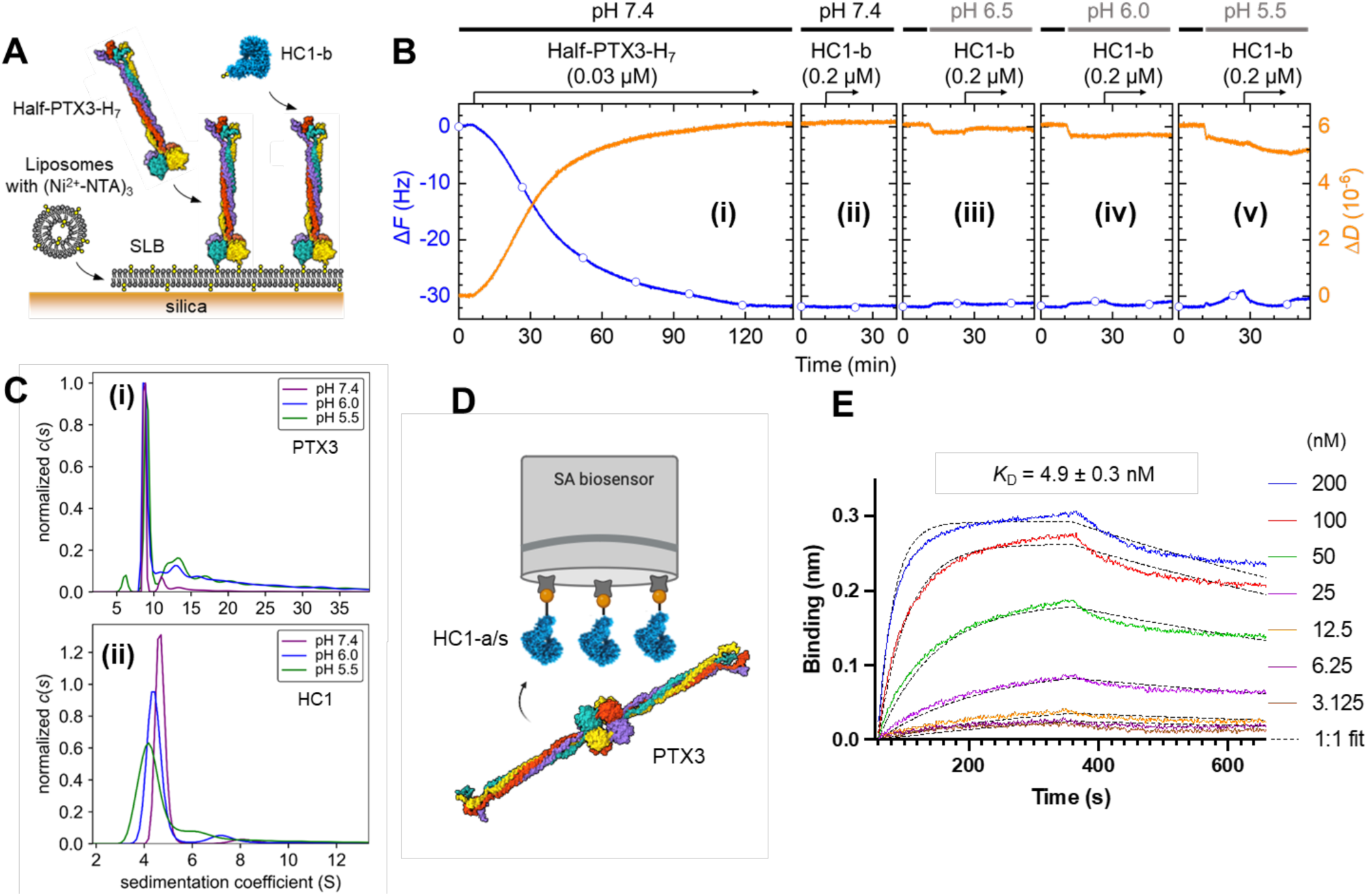
The effect of pH on the HC1-PTX3 interaction and the solution conformations of HC1 and PTX3 proteins. (**A**) Schematic of the QCM-D binding assay: Half-PTX3-H_7_ is anchored via four H_7_ loops in its C-terminal tetrameric domain to a supported lipid bilayer (SLB) made from DODA-(Ni^2+^- NTA)_3_ and DOPC lipids, allowing the N-terminal domain of PTX3 to interact with HC1 analyte in the solution phase. DOPC provides an inert background (with low non-specific binding) and the lipids are free to diffuse within the SLB, facilitating their rearrangement in plane for optimal presentation of the (Ni^2+^-NTA)_3_ moieties and multivalent attachment of Half-PTX3-H_7_ tetramers. (**B**) QCM-D responses (frequency shifts: Δ*F* (blue line with symbols); dissipation shifts: Δ*D* (orange line); data from overtone *i* = 5 are shown) upon ligand anchorage (**i**) and analyte incubations (**ii-v**). Half-PTX3-H_7_ was anchored at pH 7.4 (**i**); HC1-b incubation at different pH values (7.4 (**ii**), 6.5 (**iii**), 6.0 (**iv**) and 5.5 (**v**)) proceeded in independent experiments, where the Half-PTX3-H_7_ surface at pH 7.4 is the starting point in all cases. Arrows above the graphs represent the start and duration of protein incubations (with concentrations as indicated); during remaining times, buffer alone was flowed over the sensor surface; horizontal lines at the top indicate the solution pH (7.4: black; 6.5 to 5.5: grey). (**C**) AUC sedimentation velocity analyses for (**i**) PTX3 and (**ii**) HC1-a/s at pH 7.4 (violet), 6.5 (blue) and 5.5 (green). (**D**) Schematic of the BLI binding assay: HC1-a/s is immobilised on streptavidin-coated BLI sensors via its C-terminal Strep II tag, for interaction with PTX3 in the solution phase. (**E**) Representative BLI sensorgrams for the interactions of immobilised HC1-a/s with various PTX3 concentrations (200 nM to 3.13 nM; prepared by serial dilution and colour coded as indicated) in MBS (10 mM MES, 150 mM NaCl, pH 5.5), showing the association phase (60-360 s) and the subsequent dissociation phase (>360 s), upon rinsing in buffer. Dashed black lines are fits to the data, assuming a 1:1 interaction. The mean *K*_D_ value ± SD was derived from *n* = 3 independent experiments.

We used the quartz crystal microbalance with dissipation monitoring (QCM-D) technique to confirm proper surface functionalisation including the specific attachment of Half-PTX3-H_7_ via its H_7_ loops (Figure S9), and to monitor subsequent HC1 binding. Anchorage of PTX3 to SLBs is evident in Figure 5Bi from the decrease in the QCM-D frequency shift (Δ*F*) during PTX3 incubation. Binding remained fully stable upon subsequent rinsing in buffer at pH 7.4 (Figure 5Bi). The ratio of the dissipation shift (Δ*D*) over frequency shift (-Δ*F*) at saturation (Δ*D*/-Δ*F* = 0.2 × 10^-6^ Hz^-1^) is relatively high and indicative of a flexible conformation, consistent with our structural studies (see above), and being unperturbed by attachment to the surface. Indeed, Δ*D*/-Δ*F* ratios of 0.2 × 10^-6^ Hz^-1^ have previously been reported for membrane-bound septin proteins with flexible coiled-coil domains (Szuba et al., 2021), whereas a more globular protein such as streptavidin attains Δ*D*/-Δ*F* ratios of <0.01 × 10^-6^ Hz^-1^ when pre-absorbed on gold, and between 0.015 and 0.08 × 10^-6^ Hz^-1^ (depending on surface coverage) when linked via biotins on a short, flexible, linker to an SLB (Johannsmann et al., 2009).

As can be seen from Figure 5B, QCM-D analysis revealed no detectable interaction of recombinant biotinylated-HC1 (HC1-b; Table 2) with PTX3 at pH 7.4 (ii) or 6.5 (iii); see full description of HC1 constructs in Materials and Methods and corresponding SDS-PAGE gels in Figure S10). However, binding responses were seen as the pH was decreased to 6.0 (iv) and 5.5 (v), indicating that an acidic environment promotes HC1 binding. The QCM-D responses for the interaction of Half-PTX3-H_7_ with HC1 are relatively small (-Δ*F* < 5 Hz; at pH 5.5). For comparison, considering that HC1 is ∼11 nm in length (Briggs et al., 2020), one would expect negative frequency shifts up to 60 Hz, if HC1 was binding as a dense monolayer on a planar surface (see Materials and Methods). However, we verified that the QCM-D responses for HC1 are entirely due to specific binding to PTX3 (Figure S11). Moreover, a swap in the interaction geometry, with SLB-anchored HC1-b/s ligand (Table 2) and PTX3 as the analyte showed strong and specific binding at pH 5.5 (Figure S12). This suggests that the low responses with SLB-anchored Half-PTX3-H_7_ are likely due to film compaction, which would increase the Δ*F* upon HC1 binding (thus partially offsetting the overall decrease) and decrease Δ*D,* as is indeed the case at pH 5.5 (Figure 5Bv).

**Table 2.** Mass spectrometry analysis of HC1 constructs used in the study.

| <sup>a</sup>Construct | <sup>b</sup>Theoretical Mass (Da) | Experimental mass (Da) | Corresponding Figure number where used |
| --- | --- | --- | --- |
| H<sub>6</sub>-HC1 | 73,805.8 | 73,805.8 | S12C |
| HC1-a/s | 73218.0 | 73217.0 | 5C, 5E, 8C, S13, S14A-F |
| HC1-a | 71962.4 | 71964.0 | S11B |
| HC1-b/s | 73,444.3 | 73,444.3 | S11C, S12B |
| HC1-b | 72,189.0 | <sup>c</sup>N.D. | 5B, 7, S11A |
| ΔvWFa_HC1-a/s | 51,930.0 | 51,932.3 | 8C, S14G-H |

<sup>a</sup>Apart from H<sub>6</sub>-HC1 (Baranova et al., 2013), all other constructs were made as part of this study. <sup>b</sup>Masses for oxidised proteins containing two disulphide bonds and lacking N-terminal methionine. <sup>c</sup>N.D. = Not determined.

To understand the structural basis for this pH regulation, PTX3 and HC1-a/s were analysed by sedimentation velocity analytical ultracentrifugation (AUC), a technique that can detect changes in protein hydrodynamic size. The sedimentation coefficient of PTX3 was essentially unchanged at pH 7.4, 6.0 and 5.5 (8.8 ± 0.2 S; Figure 5Ci, Figure S13), indicating that on its own in solution PTX3 does not undergo a significant conformational change across this pH range. In contrast, a clear shift in sedimentation coefficient was measured for HC1, increasing from 4.1 S at pH 5.5 to 4.7 S at pH 7.4 (Figure 5Cii, Figure S13), indicating that HC1 becomes more compact as the pH is lowered, which is consistent with QCM-D data for immobilised HC1 (Figure S12B). However, the size distribution of sedimenting species, defined by the width of the peaks, becomes broader suggesting that HC1 also adopts a wider range of conformations at acidic pH.

Considering that the HC1 binding responses seen by QCM-D, and the shift in sedimentation coefficient determined by AUC, were largest at pH 5.5, we proceeded with further analyses of PTX3-HC1 interactions at this pH. To quantify the interaction kinetics, biolayer interferometry (BLI) was employed with HC1-a/s immobilised to a streptavidin-coated sensor via its C-terminal Strep II tag, and PTX3 as the analyte in the solution phase (Figure 5D,E and Table S3); no immobilisation was observed with the HC1-a control lacking the Strep II tag (data not shown). In this assay, the HC1-a/s surface density was kept deliberately low (∼0.4 nm binding response) to limit the stoichiometry of the interaction of (octameric) PTX3 with the immobilised HC1 ligands, whilst providing an acceptable signal-to-noise ratio. Under these conditions, the association and dissociation data (collected over a range of concentrations) could be fitted well by a 1:1 interaction model, i.e., one octamer of PTX3 per immobilised HC1 molecule (Figure 5E) leading to a *K*_D_ = 4.9 ± 0.3 nM. With higher surface densities of HC1-a/s, we found that binding curves gave a good fit with the heterogeneous analyte model but not the 1:1 binding model (data not shown), indicative of the occurrence of some multivalent interactions. These data (Figure 5E) indicate that a monovalent interaction between PTX3 and HC1 (and perhaps the other HCs) is of high affinity (at pH 5.5).

### An acidic environment promotes PTX3 binding to HC•HA complexes

We have previously found that PTX3 only binds to HC•HA matrices at pH 7.4 if it is present at the time of HC transfer onto HA (Baranova et al., 2014). Based on the results described above (Figure 5), we hypothesised that PTX3 would be able to incorporate into preformed HC•HA complexes at pH 5.5. This was tested using a previously established experimental set up (Baranova et al., 2014) with films composed of HA polysaccharides (Figure 6A). QCM-D analysis demonstrated successful binding of reducing-end biotinylated HA to a streptavidin-covered SLB, and subsequent co-incubation of TSG-6 and IαI entailed further binding responses (Figure 6Bi), consistent with HC•HA film formation, as previously reported (Baranova et al., 2014, 2013).

**Figure 6.**
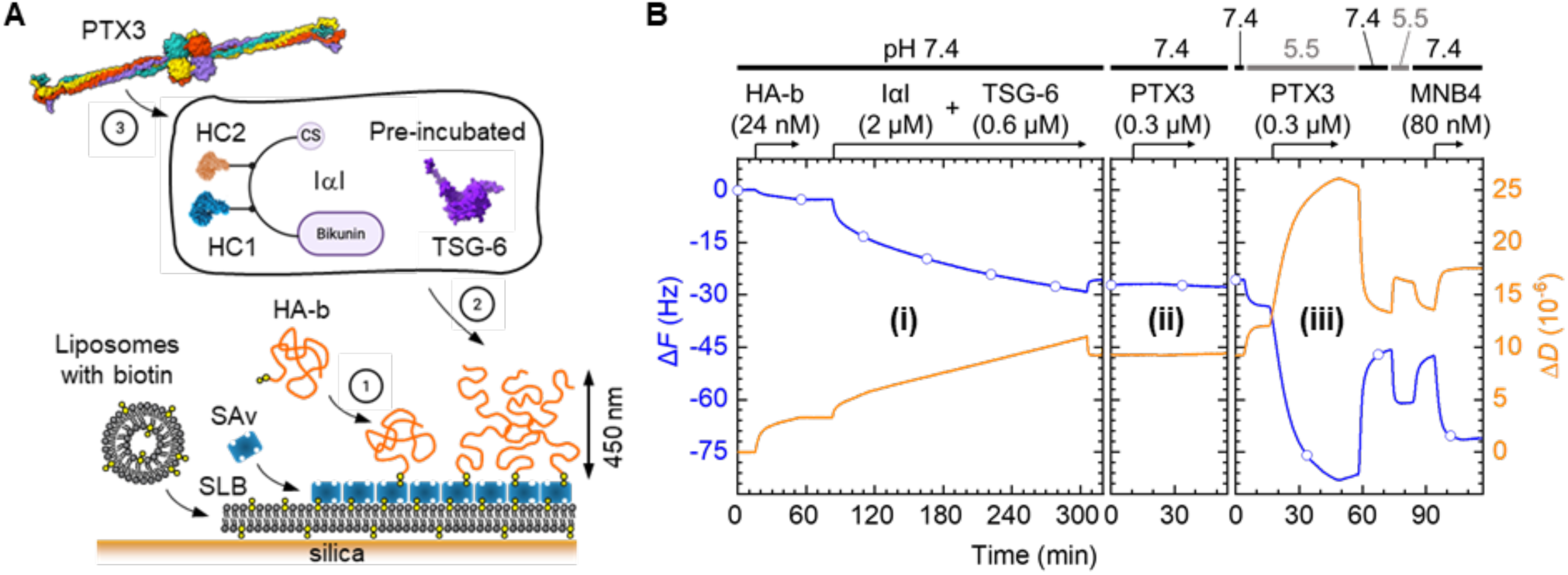
PTX3 binds to pre-formed HC•HA complexes at pH 5.5. (**A**) Schematic of the QCM-D binding assay: 1. HA polysaccharides (*M*_W_ = 840 ± 40 kDa) are anchored via a biotin tag at their reducing end (HA-b) to a streptavidin (SAv)-coated supported lipid bilayer (SLB); 2. HC•HA films are formed by co-incubation of the HA film with IαI and TSG-6 (pre-mixed for 2 h) and 3. PTX3 binding is subsequently monitored. (**B**) QCM-D responses (frequency shifts: Δ*F* (blue line with open circles); dissipation shifts: Δ*D* (orange line); data from overtone *i* = 5) upon HA anchorage and HC•HA film formation (**i**), and subsequent exposure to PTX3 at pH 7.4 (**ii**) and pH 5.5 (along with pH cycling and binding of anti-PTX3 antibody MNB4) (**iii**). Buffer conditions were HBS (10 mM HEPES, pH 7.4, 150 mM NaCl) or MBS (10 mM MES, pH 5.5, 150 mM NaCl). Streptavidin-coated SLBs were formed as shown in Figure S12; arrows above the graphs represent the start and duration of HA and protein incubations (with concentrations as indicated); during remaining times buffer alone was flowed over the sensor surface; horizontal lines at the top indicate the solution pH (pH 7.4: black; pH 5.5: grey). Data are representative of two independent experiments.

PTX3 did not bind to the preformed HC•HA films at pH 7.4 (Figure 6Bii), as observed before (Baranova et al., 2014). A reduction in pH to 5.5 had a mild impact on the HC•HA film itself (see 4 to 17 min in Figure 6Biii), but strikingly led to a strong response upon PTX3 incubation (17 to 48 min in Figure 6Biii). PTX3 remained rather stably bound upon rinsing in pH 5.5 (48 to 58 min in Figure 6Biii), and a control experiment showed that PTX3 does not bind to the SAv surface (Figure S12D), validating the specific (and substantial) interaction of PTX3 with HC•HA at pH 5.5.

Interestingly, a change from pH 5.5 to pH 7.4 following PTX3 binding (48 to 73 min in Figure 6Biii) led to the release of some but not all PTX3, accompanied by compaction and rigidification of the HA film. The pH-induced changes in film morphology were reversible, as evident from the changes in frequency and dissipation when cycling between pH 7.4 and pH 5.5 (73 to 93 min in Figure 6Biii; a Δ*F* decrease with a concomitant Δ*D* increase indicates swelling and softening, and opposite responses compaction and rigidification. Subsequent binding of the anti-PTX3 antibody MNB4 corroborated the stable retention of PTX3 in the film at pH 7.4 (93 to 108 min in Figure 6Biii).

### Stoichiometry of HC1 binding to PTX3

As noted previously, PTX3 is composed of an octamer made of 8 identical protomers (Inforzato et al., 2010, 2008), thus an important question is how many HCs can bind simultaneously to an individual PTX3 molecule. In the case of fibroblast growth factor-2 (FGF2), we previously found that there were only 2 binding sites per PTX3 octamer (Inforzato et al., 2010), whereas tetrameric forms of PTX3 can stabilise the cumulus matrix (Ievoli et al., 2011; Inforzato et al., 2008; Scarchilli et al., 2007) indicating that each TND_PTX3 is likely to be able to bind at least 2 HCs. To determine the HC1 binding stoichiometry, we deployed the Half-PTX3-H_7_ construct onto Ni^2+^-NTA-SLBs (as shown in Figure 5A) in the context of spectroscopic ellipsometry (SE), a technique that is well suited for the quantification of protein surface density (Richter et al., 2014).

To avoid any negative impact of surface crowding on HC1 binding, we kept the surface density of Half-PTX3-H_7_ relatively low: an areal mass density (*AMD*) at saturation of 26 ± 9 ng/cm^2^(exemplified in Figure 7A; at 110 min) corresponds to a molar surface density of Γ = 0.15 ± 0.05 pmol/cm^2^ (Figure 7B), and a root-mean-square distance of 35 ± 6 nm between the immobilised Half-PTX3-H_7_ molecules. At pH 5.5, when incubated at 700 nM concentration (i.e., well above the *K*_D_; see Figure 5E), HC1 bound stably to Half-PTX3 (Figure 7A; 140 to 200 min), and from the ratio of the proteins’ *AMD*s at equilibrium, and molecular masses, a binding stoichiometry of 4.0 ± 0.8 HC1 per Half-PTX3-H_7_ tetramer was obtained (*n* = 3; Figure 7C). This indicates that up to 8 HC1 molecules can bind per PTX3 octamer, providing a mechanism for the multivalent interaction of PTX3 with HC•HA complexes; i.e., with the potential to form a ‘dense’ crosslinking node that brings together multiple HA chains.

**Figure 7.**
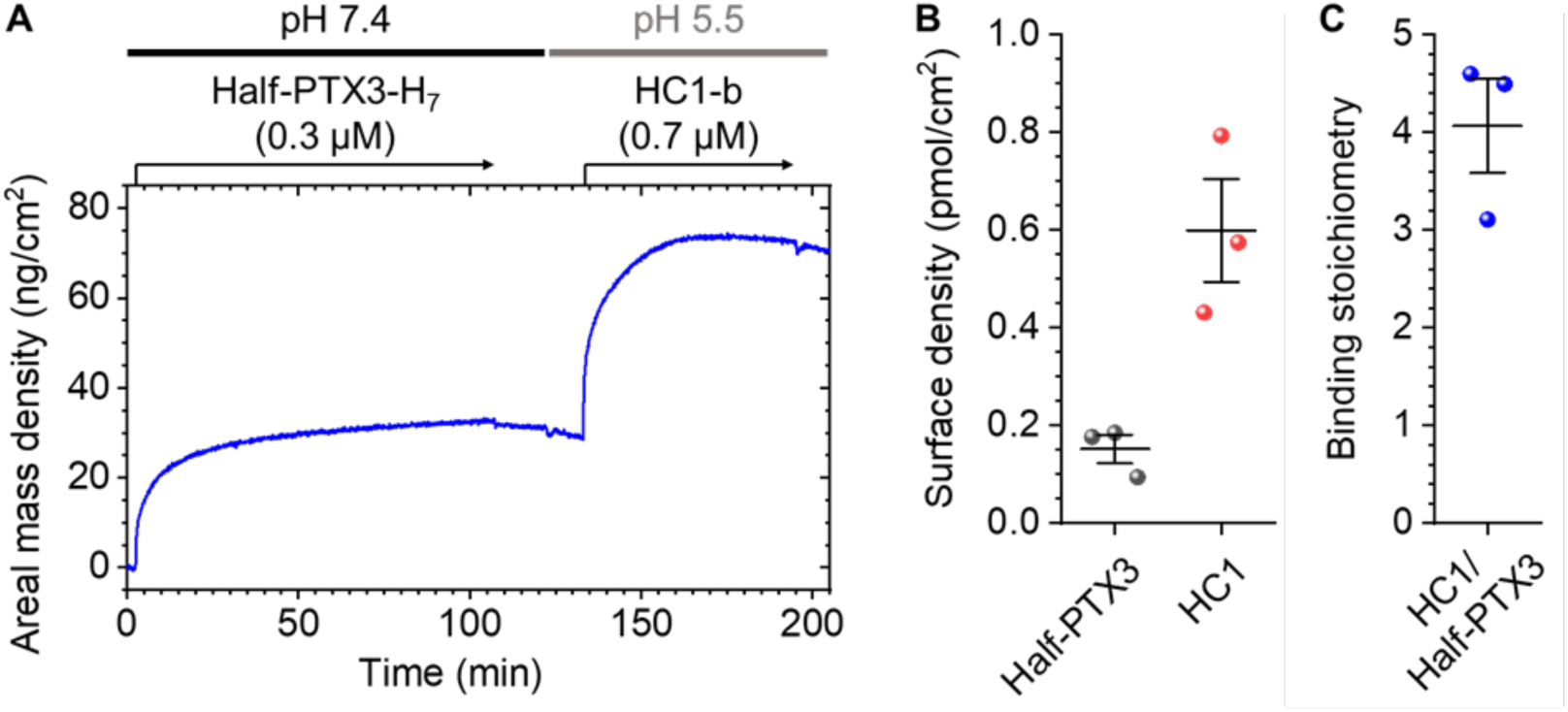
Stoichiometry of HC1 binding to PTX3. Data from spectroscopic ellipsometry (SE) for HC1-b binding to half-PTX3-H_7_ immobilised onto the biosensor surface. SE detects binding through the associated changes in the refractive index near the surface, with areal mass densities being accurate (i.e., insensitive to the organisation of the biomolecular film) up to thickness values of tens of nm. (**A**) Areal mass density vs. time for the binding of Half-PTX3-H_7_ to SLBs containing 0.4 mol-% DODA-(Ni^2+^-NTA)_3_ lipids, and subsequent binding of HC1-b. Arrows above the graph represent the start and duration of protein incubations (with concentrations as indicated); during remaining times the surface was exposed to buffer alone (either HBS (10 mM HEPES, pH 7.4, 150 mM NaCl) or MBS (10 mM MES, pH 5.5, 150 mM NaCl); horizontal lines at the top indicate the solution pH (pH 7.4: black; pH 5.5: grey). (**B-C**) Surface densities at saturation (**B**), and resulting binding stoichiometry (**C**), obtained with Half-PTX3-H_7_ (*M*_W_ = 170 kDa) and HC1-b (*M*_W_ = 72.2 kDa). In **B** and **C**, mean values ± SD are shown for *n* = 3 independent experiments.

### The N-terminal region of PTX3 is involved in the interaction with HC1

To gain insight into the location of the interaction sites on PTX3 and HC1, we used BLI with various HC1 and PTX3 constructs (Figure 8A,B; Figure S14). As can be seen from Figure 8C,D, neither removal of the von Willebrand Factor A domain from HC1 (ΔvWFA_HC1; *K*_D_ = 6.6 ± 0.3 nM) nor deletion of the first 24 amino acids from PTX3 (Δ1-24_PTX3; *K*_D_ = 7.2 ± 0.4 nM) led to an appreciable change of the interaction strength compared to the full length proteins (*K*_D_ = 4.9 ± 0.3 nM; Figure 4E) indicating that these regions of HC1 and PTX3 do not mediate the interaction.

**Figure 8.**
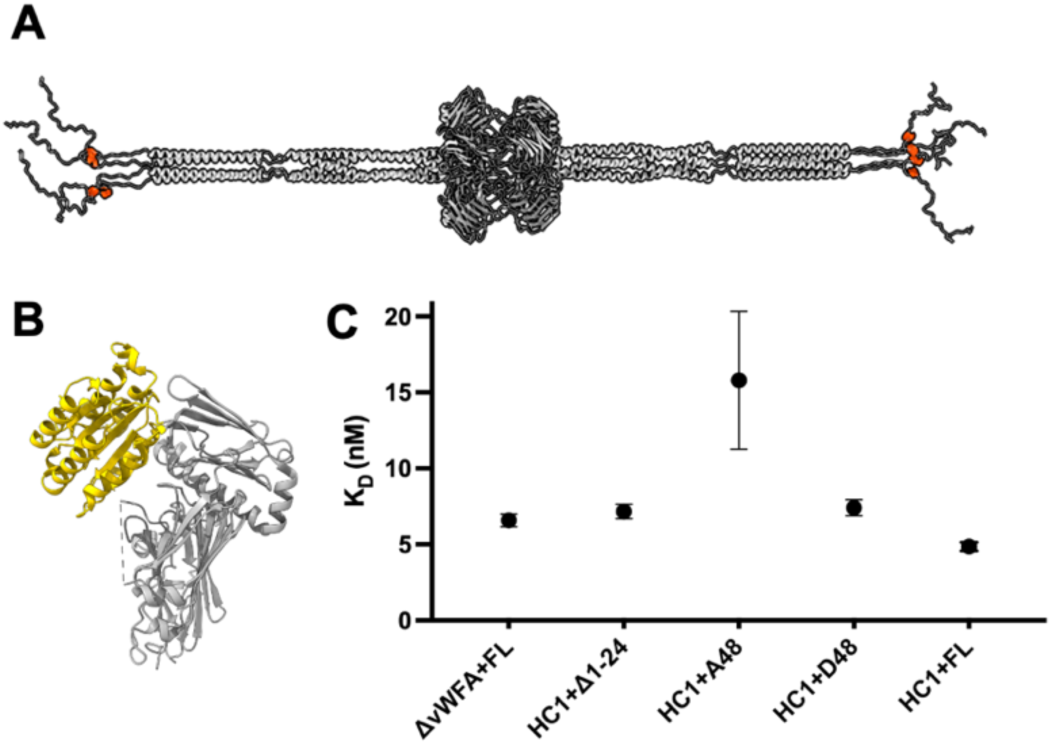
Towards identifying the HC1-PTX3 interaction sites. (**A**) PTX3 structure with the position of the polymorphic A48D residue highlighted in orange. (**B**) HC1 (6fpy.pdb) structure with the vWFA domain coloured in yellow. (**C**) Comparison of interaction affinities (*K*_D_) of different HC1 and PTX3 constructs (HC1: ΔvWFA = ΔvWFA_HC1. PTX3: Δ1-24 = Δ1-24_PTX3; A48 = A48-PTX3; D48 = D48-PTX3; FL = PTX3), as measured by BLI (mean ± SD for *n* = 3 technical repeats; see Table S3).

Previously it has been suggested that the HC binding site corresponds to residues 87-99 of PTX3 based on competition for the IαI-PTX3 interaction by the anti-PTX3 monoclonal antibody MNB4, which is reported to recognise this region (Noone et al., 2022; Scarchilli et al., 2007). Contrary to this we found that MNB4 could interact with PTX3 when already bound to HC•HA complexes (Figure 6Biii), with a substantial and monophasic binding response. The fact that the antibody binding to PTX3/HC•HA reached a clear equilibrium is indicative of a non-competitive interaction. Based on these data it seems unlikely that HC1 shares an overlapping binding site on PTX3.

Preliminary experiments where our model of TND_PTX3 and the crystal structure of HC1 were docked together using AlphaFold (data not shown) indicated that HC1 might interact with PTX3 in the ‘linear’ region (residues 47-54), i.e., between the disordered N-terminal peptide (residues 18-46) and the first coiled-coil (residues 55-100). Consistent with this prediction, constructs representing two polymorphic variants of PTX3, A48-PTX3 and D48-PTX3 (see Figure 8A for the location of amino acid 48 in the protein) (Barbati et al., 2012) both expressed in HEK Expi293F cells, were found to have different affinities to HC1 with *K*_D_ values of 15.8 ± 3.7 nM and 7.41 ± 0.4 nM, respectively (Figure 8C).

Additional BLI assays with A48-PTX3, carried out at the low HC1 surface density (∼0.4 nm binding response), showed that a reduction in the NaCl concentration (from 150 to 50 mM) enhanced binding to PTX3 whereas an increase to 250 mM NaCl virtually eliminated the interaction (Figure S14A,B). This indicates that the binding of HC1 to PTX3 involves ionic interactions, which given the pH-dependency may involve histidine residues. Furthermore, the interaction between PTX3 and the ΔvWFA_HC1 construct (that is lacking the MIDAS site) was unaffected by the presence of EDTA (Figure S14C) demonstrating that the interaction between HC1 and PTX3 does not require divalent metal ions. Previous studies reported that the binding of IαI to PTX3 was metal ion dependent (Scarchilli et al., 2007), which appears inconsistent with our findings.

Overall, the studies presented here indicate that HC1 binds via a site outside of its vWFA domain, in a salt-dependent and cation-independent manner, to a region of PTX3 towards the N-terminal of the protein, i.e., via a sequence including amino acid 48 (where the D48 allotype binds more tightly than the A48 variant). The discovery that up to eight HC1 molecules can bind to each PTX3 octamer indicates that PTX3 represents a major node in the crosslinking of HC•HA. This interaction is clearly tightly regulated only occurring at pH 7.4 when HC•HA complexes form in the presence of PTX3 but can also occur under acidic conditions such that PTX3 can incorporate into preformed HC•HA matrices at pH 5.5.

## DISCUSSION

PTX3 has a fundamentally important role in stabilising HA-rich extracellular matrices in the context of normal physiology and this activity has also been implicated in several pathological processes (Baranova et al., 2014; Brunetta et al., 2020; Inforzato et al., 2010; Massimino et al., 2023; Ogawa et al., 2017; Salustri et al., 2004; Scarchilli et al., 2007; Tseng, 2016). In addition, PTX3 is well established as a humoral fluid phase pattern recognition molecule controlling many aspects of innate immunity, including regulating the complement system in the context of cancer and infectious disease (Bonavita et al., 2015; Mantovani and Garlanda, 2023; Parente et al., 2021; Stravalaci et al., 2020). Here we have determined a structure for PTX3 based on a combination of X-ray crystallography, cryo-EM and molecular modelling and have generated new mechanistic insights into how this multimeric protein interacts with an archetypal heavy chain (HC1) of the IαI family of proteoglycans to form a major, and highly regulated, crosslinking node within the ECM.

The structure determined here for PTX3 (8PVQ.pdb) has many features in common with that recently described by Noone *et al*. (7ZL1.pdb (Noone et al., 2022)), i.e., being composed of an octamer of eight identical protomers with a central body formed of C-terminal pentraxin domains and long tetrameric coiled-coil extensions on either side of this formed of the N-terminal domains. The overall organisation is similar to what we proposed previously (Inforzato et al., 2010), albeit being symmetrical and with much longer N-terminal extensions, and is fully compatible with the experimentally determined disulphide bond network (Inforzato et al., 2008). The central cubic arrangement of 8 C_PTX3 domains from our study (denoted the OCD_PTX3), is essentially identical to that of 7ZL1.pdb, albeit informed by our high-resolution (2.4 Å) crystal structure for the PTX3 pentraxin domain.

Here we identified a network of ionic interactions between pentraxin domains that stabilise the assembly of four PTX3 protomers into a tetramer. Furthermore, salt bridges (seen in both our cryo-EM and crystal structures) connect the two tetramers into an octamer, which likely occurs prior to the formation of interchain disulphides between C317 and C318. These important observations provide novel mechanistic insights into the biosynthesis of PTX3 and the formation of oligomers that are inextricably linked with its function (Ievoli et al., 2011; Inforzato et al., 2010, 2008). Coupled with the finding that a single PTX3 molecule can interact simultaneously with up to eight heavy chains, this provides a mechanism whereby PTX3 can stabilise HC•HA containing matrices critical to physiological processes (e.g., cumulus matrix expansion prior to ovulation (Ievoli et al., 2011; Salustri et al., 2004; Scarchilli et al., 2007) and providing the amniotic membrane with potent anti-inflammatory, anti-angiogenic and anti-fibrotic properties (Tseng, 2016)).

One potential structural difference in our study from that of Noone *et al*. (Noone et al., 2022) is in the flexibility of the tetrameric N-terminal domains. In both studies two hinges are proposed at similar regions of the protein, however, Noone *et al*. (Noone et al., 2022) described a much more dramatic bending around Hinge 2 based on cryo-EM and negative staining EM data (with the N-terminal region (residues 18-102) moving by approx. 60° relative to the first coiled-coil region (residues 107-172). While we cannot rule out that such extreme flexibility occurs within some PTX3 molecules, this seems unlikely to be a general feature of the protein based on our SAXS data for the PTX3 constructs analysed in solution. This difference notwithstanding, the flexible nature of the tetrameric N-terminal domains could be important in the crosslinking of HC•HA. For example, the cumulus oocyte complex is the softest elastic tissue currently known, with a Young’s modulus of < 1 Pa (Chen et al., 2016). Given that HC•HA/PTX3 complexes are the major structural component of the cumulus matrix they must be responsible for its mechanical properties, i.e., being very soft but also very resilient. Our finding that 8 HC1 molecules can interact simultaneously with a single PTX3 octamer suggests that HC•HA/PTX3 complexes do form a major crosslinking node, linking together multiple HA chains, that likely contributes to the resilience of the matrix, especially since the affinity of HC1 for PTX3 was found here to be in the low nM range. When combined with weaker binding processes, as described previously for HC1-HC1 homotypic dimers (Briggs et al., 2020), this could engender the elasticity of the COC. Clearly, further work is needed to determine the affinities for the other HC-PTX3 and HC-HC interactions to obtain a fuller understanding of the mechanisms of crosslinking.

We have found that the D48 allotype of PTX3 binds to HC1 with a 2-fold higher affinity compared to A48-PTX3. This provides evidence that this amino acid forms part of the HC1-binding site and that this is located towards the N-termini of PTX3. This is a structurally unresolved part in our model, where residue 48 sits between the two cysteines (C47 and C49) that form interchain disulphide bonds (Inforzato et al., 2008), so it is possible that this region has a defined structure. However, given we do not know precisely how the four chains in a tetramer are linked by the C47/C49 disulphides or indeed where on HC1 the binding site for PTX3 is located (except that it is not within the vWFA domain) it is not currently possible to model this interaction.

What is apparent is that the binding of HC1 close to the N-terminal ends of PTX3 means that a large distance can be spanned when PTX3 links together two HC1•HA complexes, i.e., ∼450 Å. This combined with the flexibility of the PTX3 protein (see above), might explain why only a relatively small contraction in HA film thickness (and little rigidification) occurs on the incorporation of PTX3 into an HC•HA network (Baranova et al., 2014). In this regard, based on the large size of this crosslink we would predict that HA modified with HCs and PTX3 would retain many of its solution properties, and notably, a very open structure as required to make an ultra-soft matrix (Chen et al., 2016). However, it is possible that in contexts where HA is heavily decorated with HCs this would lead to a much greater degree of crosslinking and rigidification, i.e., given our finding that PTX3 can interact simultaneously with up to eight HC1 molecules. It is conceivable that in the tissue context each PTX3 molecular features a variable number of HA-attached HCs, either in close proximity (i.e., on the same N-terminal arm) or spaced far apart (i.e., on opposing N-terminal arms), leading to HA matrices with a spectrum of mechanical properties and functions. Moreover, whether the other HCs can bind to the same binding site as HC1, or whether they interact at other locations on PTX3, is not yet described, but this could also have an important bearing on what crosslinks are formed, e.g., depending on the HC•HA composition.

The A48 and D48 alleles are both common in the European population with a frequency of 42% vs. 58%, respectively (gnomAD variant ID: 3-157155314-C-A; see www.gnomAD.broadinstitute.org [gnomad.broadinstitute.org). This polymorphism is part of a haplotype that has been associated with the occurrence of dizygotic twins in mothers from The Gambia, where the A allele (coding for D48) is more common than the C allele (coding for A48) amongst twinning mothers (Sirugo et al., 2012). Our finding that D48-PTX3 binds with greater affinity to HC1 than A48-PTX3, indicates that this allotype may be incorporated into the cumulus matrix at a higher concentration. Given that PTX3 has been suggested to play a role in sperm attachment to the COC (Salustri et al., 2004) it is feasible that this could contribute to more frequent fertilisation resulting in a larger number of fraternal twins being conceived.

Interestingly, the *PTX3* polymorphisms (including the A48D exonic variant) have also been linked to protection from pulmonary tuberculosis in Guinea-Bissau (Olesen et al., 2007), *P. aeruginosa* airway infection in European patients with cystic fibrosis (Chiarini et al., 2010) and invasive pulmonary aspergillosis in diverse clinical settings in Caucasians (Parente et al., 2021). Our findings might provide a mechanistic explanation for these observations given that the formation of HC•HA complexes have been documented in other diseases of the airway (Bell et al., 2019; Queisser et al., 2021; Tang et al., 2024), i.e., greater binding of the D48 isoform to HC•HA may increase the level of PTX3 within lung extracellular matrices thereby enhancing PTX3-mediated regulation of the innate immune system and improving clearance of these pathogens.

The position of the HC1-binding site that we have proposed (based on data obtained with our A48-PTX3 and D48-PTX3 constructs) is different from that indicated by studies where the interaction of IαI with PTX3 is inhibited by the MNB4 monoclonal antibody (Scarchilli et al., 2007), the epitope of which has been mapped to amino acids 87-99 (Camozzi et al., 2006). In our study the addition of MNB4 did not affect the binding of PTX3 to HC•HA and, moreover, we found that the interaction of HC1 with PTX3 was metal ion-independent, while the binding of IαI to PTX3 requires Mg^2+^ ions (Scarchilli et al., 2007); there is no known metal ion binding site in PTX3 whereas Mg^2+^ is accommodated in the MIDAS of HC1 (Briggs et al., 2020). Furthermore, we found that the interaction between HC1 and PTX3 was not affected by the removal of the vWFA domain, where the MIDAS is located. Thus, there are many apparent contradictions between our data and the previous studies (Scarchilli et al., 2007). One intriguing possibility is that the binding of HC1 to PTX3 is distinct to that of IαI (i.e., with regard to the position where they bind and the nature of the interaction). As described above, PTX3 only incorporates into the HC•HA matrix at pH 7.4 if PTX3 and IαI interact prior to the formation of HC•HA. Perhaps, this interaction is metal ion-dependent and maps to the 87-99 region. Further studies will be needed to test this.

Regardless of the precise details, it is clear that the PTX3-mediated crosslinking of HC•HA is tightly regulated at pH 7.4 (Baranova et al., 2014). Consistent with our previous data, we found here that PTX3 can neither bind to free HC1 nor incorporate into preformed HC•HA complexes at this pH value. This is suggestive that only under very precise conditions can a HA matrix crosslinked by HC•HA/PTX3 complexes be formed (e.g., when HA, PTX3 and TSG-6 are expressed by cumulus cells and IαI ingresses into the ovarian follicle (from serum) in the preovulatory period (Carrette et al., 2001; Hess et al., 1998; Mukhopadhyay et al., 2001; Salustri et al., 2004)). Importantly, however, we found that PTX3 can bind to HC1 at pH 6.0 and more tightly at pH 5.5 and incorporates strongly into a HC•HA film at pH 5.5. HC1 (but not PTX3) was found to undergo a conformational change between pH 7.4 and 5.5, where this may be necessary for the interaction to occur (e.g., unmasking a binding site); previously we have reported that there is a change in HC1 conformation when it forms a homotypic interaction (Briggs et al., 2020). It is possible, therefore, that the change in conformation of HC1 with pH mimics a ligand induced-conformational change that occurs in the context of the IαI-PTX3 interaction at pH 7.4.

Our current studies reveal that under conditions of tissue acidosis PTX3 could contribute to the stabilisation of HA matrices where HC modification has previously occurred (i.e., sites of prior inflammation). For example, pH could regulate HA crosslinking in lung pathologies (e.g., asthma and viral infections), where HC•HA formation, PTX3 and acidosis have all been implicated (Bell et al., 2019; Brunetta et al., 2020; Forteza et al., 2007; Koussih et al., 2021; Lauer et al., 2015; Queisser et al., 2021; Tang et al., 2024); e.g., during acute asthma a pH of 5.2 has been reported in the lower airways (Hunt et al., 2000; Ricciardolo et al., 2004). In addition, pH values as low as pH 5.6 have been reported in solid tumours (see (Erra Díaz et al., 2018)) where PTX3 is often unregulated (Bogdan et al., 2022; Chang et al., 2021) and for which a protective role for HC•HA has been identified in breast cancer (Zhang et al., 2021). In these situations, PTX3-medited crosslinking of HA would be expected to be highly adhesive to leukocytes (Zhuo et al., 2006), however, whether this is protective or drives the pathology needs to be further investigated. There is also likely to be a drop in the pH of follicular fluid following the gonadotrophin surge since this increases the concentration of inflammatory cytokines (Adamczak et al., 2021). Thus, it is possible that pH could also serve to facilitate the incorporation of PTX3 into the cumulus matrix at sites where HC•HA have already formed.

In this study, our structural work on human PTX3 and the in-depth analysis of its interaction with HC1 of the inter-α-inhibitor family of proteoglycans has generated novel insights into the mechanism by which HC•HA become crosslinked and how this is regulated. This provides an important step towards a greater understanding of the structure/function interrelationships of HC•HA/PTX3 complexes and how they contribute to the organisation of extracellular matrix in physiological and pathological processes.

## MATERIALS AND METHODS

### Protein expression and purification

#### Production of recombinant PTX3 variants

##### Plasmid and cell lines

Full-length human PTX3 (PTX3) and the C-terminal pentraxin domain (C_PTX3; see Figure S1) were expressed in CHO 3.5 cells as described previously (Bottazzi et al., 1997; Inforzato et al., 2010); the cell-line was stably transfected with pSG5 vectors (Stratagene, La Jolla, CA) encoding the 18-381 and 178-381 amino acid sequences of the human preprotein (P26022), respectively, with the latter containing 3 non-authentic amino acids (ELE) at its N-terminal end.

Additional constructs of PTX3, for HEK Expi293F cell expression, were generated by GeneArt (Themo Fisher Scientific) within the pcDNA 3.4 TOPO vector. These include: A48 and D48 polymorphic variants of PTX3 (denoted: A48-PTX3 and D48-PTX3, respectively); a C317S/C318S mutant of PTX3 (Half-PTX3); a mutant where the 316-GCCVGGG-322 sequence was replaced by HHHHHHH in PTX3 (Half-PTX3-H_7_); and a PTX3 truncation mutant with residues 1-24 of the mature protein (18-41 in P26022) deleted, which contains 2 non-authentic amino acids (EL) at its N-terminal end (Δ1-24_PTX3).

It should be noted that the PTX3 protein (made in CHO cells) corresponded to the D48 allotype while the Half-PTX3, Half-PTX3-H_7_ and Δ1-24_PTX3 constructs all had A at position 48.

##### CHO culture

CHO 3.5 cells expressing the PTX3 and C_PTX3 proteins were grown as adherent cultures in Dulbecco’s Modified Eagle’s Medium (DMEM) (Sigma-Aldrich) supplemented with 10% (v/v) heat inactivated foetal bovine serum (FBS; Sigma-Aldrich) containing 100 nM non-essential amino acids, 1 mM sodium pyruvate, 2 mM glutamine (all from Lonza) and 800 µg/mL of gentamicin G418 (Sigma-Aldrich) at 37 °C in a humidified atmosphere containing 5% (v/v) CO_2_. Conditioned medium was collected from confluent cells after 3 days (for PTX3) or 7 days (for C_PTX3) of culturing in the absence of FBS; the proteins were analysed by SDS-PAGE (Figure S2).

##### HEK EXPI culture

HEK Expi293F human cells were transiently transfected using the ExpiFectamine Transfection kit (ThermoFisher) to express the Half-PTX3, Half-PTX3-H_7_, Δ1-24_PTX3, A48-PTX3 and D48-PTX3 constructs. Cells were grown as suspension cultures at 37 °C in 125 mL Corning Erlenmeyer shaker flasks (Sigma-Aldrich) at a cell density of 3 to 5 × 10^6^ cells/mL in 30 mL of Expi293 Expression Medium (ThermoFisher) in a humidified atmosphere containing 8% CO_2_ and shaking at 125 rpm; cells were sub-cultured a minimum of 3 to 5 times before transfection at a cell density of 7.5 × 10^7^ cells/mL, according to the manufacturer’s instructions. Conditioned medium was harvested 5 to 7 days post transfection.

##### PTX3 purification

All PTX3 proteins were purified from conditioned media by a combination of immunoaffinity (IAC) and size exclusion (SEC) chromatography. IAC was performed on HiTrap NHS-activated HP 5-mL columns (GE Healthcare) coupled with 2 to 3 mg/mL of either MNB4 (an anti-PTX3 monoclonal antibody recognising an epitope in the N-terminal domain that was used to purify PTX3, Half-PTX3, Half-PTX3-H_7,_ Δ1-24_PTX3, A48-PTX3 and D48-PTX3; (Camozzi et al., 2006) or MNB1 (an anti-PTX3 monoclonal antibody that binds the C-terminal domain of PTX3 and was used to purify C_PTX3; (Camozzi et al., 2006). The columns were equilibrated with PBS at 5 mL/min, and bound proteins eluted with 100 mM glycine, pH 2.7; the eluate was monitored at A280 nm, and protein-containing fractions immediately neutralized to pH 7.4 by addition of 1 M Tris-HCl, pH 8.8. There fractions were pooled and concentrated using Vivaspin 6 (10 kDa MWCO; Sartorius) and then loaded onto Superose 6 10/300 GL (for PTX3, Half-PTX3, Half-PTX3-H_7_, Δ1-24_PTX3, A48-PTX3 and D48-PTX3) or Superdex 200 10/300 GL (for C_PTX3) SEC columns (GE Healthcare), equilibrated/run in PBS at 0.5 mL/min, with protein elution monitored at A280 nm. Protein concentration was determined by UV absorbance at 280 nm, whereby a 1 mg/mL solution of PTX3 has an A280 nm of 1.5 (corresponding to an extinction coefficient in water of 60,000 M^-1^ cm^-1^). The purified PTX3 proteins were analysed by SDS-PAGE on either NuPAGE 4-12% Bis-Tris polyacrylamide gels (under reducing conditions) or NuPAGE 3-8% Tris-acetate gels (under non-reducing conditions); see Figure S2.

#### Production of recombinant HC1 variants

A construct encoding WT HC1 protein (residues 35-672 in P19827), which has an N-terminal sequence MA<u>HHHHHH</u>VGTGSNDDDDKSPDP that includes a 6-histidine tag, prior to the first amino acid (residue 35) of the mature protein and denoted H_6_-HC1, was expressed in *E. coli* and purified as described previously (Baranova et al., 2013). This and a related construct, lacking residues 288-478 (denoted H_6_-HC1_ΔvWFA (Briggs et al., 2020)) were mutated (by Genscript) to remove the non-authentic N-terminal sequence (i.e., so that residue 35 follows the initiating methionine) and residues 653-672 (which were not resolved in the HC1 crystal structure (Briggs et al., 2020)), followed by addition of an Avi/StrepII tag (GLNDIFEAQKIEWHEGGENLYFQGSAWSHPQFEK) to the C-terminal end; these new constructs are denoted HC1-a/s and ΔvWFA_HC1-a/s, respectively.

Following expression (using the method described in (Baranova et al., 2013) cell lysates containing HC1-a/s and ΔvWFA_HC1-a/s were initially captured using Strep-Tactin Superflow resin (IBA Lifesciences) and eluted with desthiobiotin (Sigma Aldrich). Where required, the Avi-tag (GLNDIFEAQKIEWHE) of HC1-a/s was biotinylated using biotin-protein ligase (BirA; Avidity) to generate HC1-b/s. Moreover, the StrepII tag (WSHPQFEK) could be removed enzymatically by cleavage between the Q and G residues in the TEV recognition sequence (ENLYFQG) allowing generation of Avi-tagged proteins with or without a biotin (denoted HC1-b or HC1-a, respectively). A final SEC purification step was performed on a Superdex-200 column and the protein assayed by SDS-PAGE (Figure S11) and liquid chromatography mass spectrometry (LC-MS); experimental values for the constructs are presented in Table 2, which are all within 2.5 Da of their theoretical molecular masses.

### Crystallisation and crystal structure determination

#### Preparation of C_PTX3 for crystallisation

The C_PTX3 protein was purified as described previously (Bottazzi et al., 1997; Inforzato et al., 2010) and buffer exchanged from PBS into 100 mM Tris-HCl, pH 8.00 on a Superdex 200 10/300 GL column (GE Healthcare). Protein-containing fractions were pooled and concentrated to 2-3 mg/mL prior to precipitation of C_PTX3 by addition of a saturated ammonium sulfate solution in 100 mM Tris-HCl, pH 8.00 (to 60% saturation) with continuous stirring at 4 °C followed by centrifugation at 4 °C for 30 min at 16.000 × *g*. The protein precipitate was stored at −80 °C until use in crystallisation trials. A 2 mg aliquot of C_PTX3 as an ammonium sulphate precipitate was solubilised in 100 mM Tris-HCl, pH 8.00 and purified by SEC on a Superdex 200 10/300 GL column (GE Healthcare) equilibrated in the same buffer, and the eluted protein directly utilised for crystallographic screening. Drops of 400 nL (200 nL protein + 200 nL reservoir) were set up in 3 lens crystallisation plates (SWISSCI) against a suite of 6 commercial crystallisation screens (Molecular Dimensions), with plates being incubated at 4 °C. No crystals were observed over a 3-month period, however, following > 12 months a single crystal was observed to have grown from a reservoir condition comprising 10% (w/v) PEG 8000, 20% (v/v) ethylene glycol, 0.03 M of sodium fluoride, sodium bromide, sodium iodide [Morpheus B6] in 0.1 M MOPS/HEPES-Na pH 7.5. The crystal was harvested and cryo cooled in liquid N_2_ prior to X-ray data collection at Diamond Light Source beamline I04-1 (100 K, 360° sweep, 0.2° oscillation). Data were integrated, scaled, and merged using the automated xia2dials pipeline through ISPyB. The structure was determined by molecular replacement in Phaser using a search model derived from 3KQR.pdb. Subsequent cycles of iterative model building, and refinement, were carried out in COOT (Emsley and Cowtan, 2004) and PHENIX (Adams et al., 2010). Validation with Molprobity (Williams et al., 2018) and PDB-REDO (van Beusekom et al., 2018) were integrated into the iterative rebuild and refinement process. A paired refinement procedure was used to determine the final resolution cut off of 2.43 Å. Complete data collection and refinement statistics are presented in Table 1 and at 8PVQ.pdb.

### Cryo electron microscopy (cryo-EM)

#### Grid preparation

UltrAuFoil R2/2, 200 mesh surface grids (Quantifoil) were used with a 30 s glow discharge at 25 mA (Quorum Emitech K100X) to clean and charge the gold surface. 3 µL of purified PTX3 at 0.25 mg/mL were applied to the grid and blotted for 4 s at 4 °C, 100% humidity in a MarkIV Vitrobot (Thermo Fisher Scientific), before plunge freezing in liquid ethane.

#### Data acquisition

Our early studies on PTX3 in 2018/2019 (from CHO cells) using a Titan Krios 2 microscope with Falcon 4 detector (University of Leeds) generated a high resolution (3.3 Å) cryo-EM structure for the OCD_PTX3 (M.P. Lockhart-Cairns, C.W. Levy, A. Inforzato, J. Fontana, R.P. Richter and A.J. Day, unpublished data). However, the PTX3 when vitrified for cryo-EM using Quantifoil or C-flat grids suffered from preferential orientation, likely caused by the dissociation of the tetrameric N-terminal domains (TND) at the air-water interface (i.e., from inspection of 2D classes; M.P. Lockhart-Cairns, C.W. Levy, A. Inforzato, J. Fontana, R.P. Richter and A.J. Day, unpublished data). This was partially overcome by taking images at 20° tilt that presented rarer ‘side views’ of PTX3 and, in addition, we explored various surfactants, glow discharge methods, grid types and grid preparation devices (Kontziampasis et al., 2019; Levitz et al., 2022). Various datasets were also collected under the optimised UltraAuFoil grid conditions, demonstrating 5-10% full side views of PTX3 with sufficient orientation distribution. However, despite this, a structure of the TND_PTX3 could not be determined due to poor contrast.

The use of a Titan Krios microscope equipped with a Falcon 4 and Selectris energy filter, at the UK National Electron Bio-Imaging Centre (eBIC), improved the contrast sufficiently to enable resolution of electron density within the TND_PTX3 (denoted Dataset 1); see Table S2 for the acquisition parameters. A separate dataset was also collected, under the same grid conditions, on the Titan Krios 2 microscope with Falcon 4 detector (University of Leeds). This dataset (denoted Dataset 2; Table S2) did not enable us to gain the resolution on the TND_PTX3 obtained with Dataset 1, however we were able to resolve more of the TND_PTX3, with Dataset 2 being used for our 3DVA analysis.

#### Data processing

The cryo-EM datasets were processed using the CryoSPARC processing package (Kucukelbir et al., 2014; Punjani et al., 2017).

Dataset 1. The recorded movies were motion corrected on the fly using motioncorr embedded in Relion (Zheng et al., 2017; Zivanov et al., 2018). All subsequent data processing was completed in CryoSPARC. Initially, 534,468 particles were selected using the CryoSPARC blob picker to generate 2D references for template picking. Iterative rounds of template picking and 2D classification yielded particle stacks that enabled reconstruction of an *ab-initio* model that was refined in D4 symmetry using the cryoSPARC non-uniform refinement algorithm. An initial stack of 106,784 particles from a subset of 500 micrographs was extracted and classified, and 11 classes containing 22,887 particles was used to train a Topaz picking model (Bepler et al., 2019) using the RESNET8 architecture. Several iterations of topaz picking, 2D classification, and model re-training were performed across the entire dataset, including the training of models that specifically targeted side views. Several strategies were employed to obtain high quality reconstructions, particularly with respect to the coiled coil regions. These included symmetry expansion, and local refinement with various masks. The most successful technique involved stringent selection of only those high resolution 2D classes (better than 8 Å) where an ordered section of coiled coil could be visually identified. This resulted in a final stack of 23,457 particles, and the final map was generated via non-uniform refinement in D4 symmetry (see Table S4).

Dataset 2. The recorded movies were motion and CTF corrected using CryoSPARC live. All subsequent data processing was completed in CryoSPARC. The automated blob picker from CryoSPARC failed to pick side view particles, picking instead the electron dense OCD_PTX3 domain. Therefore, to ensure inclusion of representative side view orientations, we manually picked particles which had visual density for the TND_PTX3 regions. Template-based picking resulted in a total of 1,076,003 particles (including both side and ‘top’ views) that were submitted to multiple rounds of 2D classification and homogeneous refinement (with D4 symmetry applied). The refined particles were Fourier cropped from 432 pixels by four-fold for faster subsequent heterogeneous refinement steps. The box size was then restored to 432 pixels. Of the resultant 846,072 particles, ∼3% presented side views of PTX3. Using the best particles, homogeneous and non-uniform refinement was used to refine the D4 maps. A resolution of 2.7 Å was achieved for OCD_PTX3 (not shown). To extend the map further and resolve the interface between the C- and N-terminal domains, particles corresponding to the side view 2D classes were selected for homogeneous refinement with C1 symmetry. Local refinement on 14,376 as then used to achieve a higher resolution and gain more detail for the TND_PTX3 with either C1 or D4 symmetry, followed by 3DVA (Punjani and Fleet, 2021).

### Modelling of the cryo-EM structure

Molecular modelling to generate the PTX3 cryo-EM structure for residues 150-381 was performed using initial docking of the crystal structure into the cryo-EM density using Chimera (Pettersen et al., 2004). This was followed by inspection in COOT and manual rebuilding where necessary. Here refinement of atomic coordinates (real space) and B-factors was performed using PHENIX, with model quality validation performed using Molprobity. Initial refinement was performed using a dimeric structure consisting of two pentraxin domains (analogous to the crystal structure unit cell) to assist in the modelling/refinement of C317 and C317. In this regard, interchain C317-C318 disulphide bonds were formed as described in Noone *et al*. (Noone et al., 2022). This is consistent with results from disulphide bond mapping described before (Inforzato et al., 2008). However, it should be noted that while these cysteines in some protomers can form intrachain disulphides only interchain disulphide bonds were modelled here. Prior to final refinement, the full assembly was then generated by replication around a 4-fold symmetry axis parallel to the coiled-coil. It appears likely that the carboxyl group of the C-terminal residue S381 (that was not fully resolved in cryo-EM map, albeit with density present) forms a (transient) salt bridge with the sidechain of R172 from an adjacent protomer. Therefore, S381 was modelled into this position.

### Modelling of the TND region of PTX3

The structure of residues 18-149 from TND_PTX3 were modelled in order to generate a complete model for PTX3. Here the structure of 18-174 were generated using Alphafold2 (Jumper et al., 2021), with multimer settings, followed by relaxation in Amber. The predicted structures for four protomer chains were then aligned with the overlapping regions of the cryo-EM structure (residues 150-174) using PHENIX and merged with the junction at residue 153; this allowed some minor real-space refinement into experimental density using COOT with geometry restraints ensuring that the bond lengths and angles at the junction were appropriate. The combination of our cryo-EM structure and the AlphaFold prediction generated two regions of tetrameric coiled-coil (residues 55-100 and 107-172), separated by a short non-helical sequence (residues 101-106). Since C103 is known to form interchain disulphide bonds (Inforzato et al., 2008), these were modelled so as to connect two pairs of protomers. The N-terminal region of PTX3 (residues 18-54) was predicted to be unstructured, therefore, residues 47-54 were initially positioned in an extended conformation, allowing C47 and C49 to form interchain disulphides (C47-C47 and C49-C49; consistent with (Inforzato et al., 2008)). Residues 18-46 were then relaxed using the ‘Model/Refine Loops’ tool in Chimera (Modeller) (Webb and Sali, 2016) to generate the final model for PTX3 shown in Figure 2F. Here, each chain was relaxed individually, and five relaxed models were generated for each. The model that gave the best zDOPE (normalised Discrete Optimised Protein Energy) score was chosen for the final PTX3 model.

### Modelling of Half-PTX3 and Δ1-24_PTX3

From the PTX3 model described above, additional models were made for the Half-PTX3 and Δ1-24_PTX3 constructs. For the former, residues C317 and C318 were mutated to serines generating just a single tetramer comprised of four protomers. In the latter the N-terminal 24 residues were removed.

### Glycan modelling

The glycosylation patterns for the CHO and HEK Expi293F expressed PTX3 constructs differ (Figure S7). For PTX3 from CHO cells, the core-fucosylated, di-sialylated and bi-antennary glycan was of highest abundance of the five glycans identified by mass spectroscopic analysis, and was, therefore, chosen for modelling. For models of Half-PTX3 and Δ1-24_PTX3 which were expressed in HEK Expi293F cells, a core-fucosylated, tri-sialylated and tri-antennary glycan was chosen since this was most abundant (Figure S7). The respective glycan structures from the Glyprot database were attached to the models for the various constructs (described above) using AllosMod, with the glycan ‘flexed’ to give the rotamer with the best fit to the SAXS curves, compared in Allosmod-FoXS (Schneidman-Duhovny et al., 2013, 2010; Weinkam et al., 2012).

### SEC-SAXS of PTX3 proteins

SAXS data (Table S1) for three PTX3 constructs (PTX3 (6 mg/ml), Half-PTX3 (5.3 mg/ml) and Δ1-24_PTX3 (8.5 mg/ml)) were collected on beamline B21 at Diamond Light Source. The samples (45 µL) were loaded onto a Superose 6 Increase SEC column (Cytiva) equilibrated and run in TBS for PTX3, and PBS for the other constructs, at a flow rate of 0.075 mL/min (20 °C). The eluent was passed through the SAXS beamline (wavelength 0.95 Å) with 1 s exposures per frame over 620 frames, and data were collected on an Eiger x 4M detector (Dectris) at a sample-to-detector distance of 2.7 m; at this distance, scattering data resulting from objects larger than 50 nm are not recorded. Raw SAXS data images were processed using Diamond Light Source’s DAWN processing pipeline to produce normalised, integrated 1-D unsubtracted SAXS curves. The raw frames were selected for buffer subtraction using ScÅtterIV. The resulting curves were then analysed in ScÅtterIV, selecting the corresponding buffer and sample peaks.

### Biolayer interferometry of HC1-PTX3 interactions

Biolayer interferometry (BLI) binding experiments were performed on an OctetRED96 system (ForteBio) in solid-black 96-well plates at 35 °C with an agitation speed of 1000 rpm. High Precision Streptavidin Biosensors (Satorius) were hydrated in 20 mM HEPES, 150 mM NaCl, pH 7.4, 0.05% (*v*/*v*) Tween-20 for 10 minutes and the surfaces then treated with regeneration buffer (0.1 M glycine, 1 M NaCl, 1 mM EDTA, 0.1% Tween-20, pH 9.5). Following washing (in 20 mM HEPES, 150 mM NaCl, pH 7.4, 0.05% (*v*/*v*) Tween-20), HC1-a/s (25 µg/mL; ‘ligand’) was bound onto the biosensor via its StrepII tag. The loaded biosensors were then dipped into interaction buffer (20 mM MES, 150 mM NaCl, pH 5.5, 0.05% (*v*/*v*) Tween-20 buffer, unless otherwise stated) and subsequently incubated with serial dilutions of PTX3 ‘analyte’ in a 96-well plate (one row of a the 96 well plate per concentration, plus a reference row (R_ref_) without analyte; see BLI Figure legends for concentrations used). Experiments were performed as technical triplicates (one column of the 96 well plate (8 wells) per replicate, plus a reference column (C_ref_) without ligand). The biosensors were then regenerated with regeneration buffer. The data were double reference subtracted: against R_ref_ to correct for solution effects that are introduced by the buffer, and against C_ref_ to correct for non-specific binding of analyte to the sensor surface. However, no non-specific binding of analyte was observed to the Streptavidin Biosensors. The binding kinetics were determined using Octet software version 7 (ForteBio) from 3 independent experiments (Table S3).

### Formation of supported lipid bilayers for binding studies

Supported lipid bilayers were formed by the method of vesicle spreading, using small unilamellar vesicles (SUVs) with desired lipid compositions. Biotin-presenting SLBs were formed in HEPES buffered saline (HBS; 10 mM HEPES, pH 7.4, 150 mM NaCl) and used to support a monolayer of streptavidin for anchorage of proteins (with biotin and/or StrepII tags) and hyaluronan (with a biotin tag at the reducing end). Ni^2+^-NTA presenting SLBs were formed in HBS supplemented with 10 mM NiCl_2_ (to load the NTA moieties with Ni^2+^ and to facilitate SLB formation) and used to anchor proteins with a polyhistidine tag. DOPC lipids served as a background to dilute lipid analogues with headgroups carrying either a biotin or a (Ni^2+^-NTA)_3_ moiety.

SUVs were prepared by sonication using established procedures (Richter et al., 2003). SUVs contained either 5 mol-% DOPE-cap-biotin (Avanti Polar Lipids) or 0.4 to 0.5 mol-% DODA-(Ni^2+^-NTA)_3_ in a background of DOPC (Avanti Polar Lipids) in HBS. DODA-(Ni^2+^-NTA)_3_ is a lipid analogue with two oleoyl tails and a chelator headgroup comprising three nitrilotriacetic acid moieties, which was prepared as described earlier (Beutel et al., 2014) and kindly provided by Changjiang You and Jacob Piehler (Osnabrück University, Germany). The relatively high DOPE-cap-biotin content in SLBs enabled the formation of a dense monolayer of streptavidin (Migliorini et al., 2014). In contrast, the relatively low content of DODA-(Ni^2+^-NTA)_3_ was selected to limit the surface density of half-PTX3-H_7_, thus avoiding steric hindrance for HC1 binding (Sobrinos-Sanguino et al., 2019; Zahn et al., 2016).

### Quartz crystal microbalance with dissipation monitoring (QCM-D) to probe PTX3 interactions with HC1 and HC•HA complexes

QCM-D measurements were performed with a Q-Sense E4 system equipped with Flow Modules (Biolin Scientific) on silica-coated sensors (QSX303; Biolin Scientific). The flow rate was controlled with a syringe pump (Legato; World Precision Instruments) at typically 20 µL/min. The working temperature was 23 °C. Before use, sensors were treated with UV/ozone for 30 min. QCM-D data were collected at six overtones (*i* = 3, 5, 7, 9, 11 and 13, corresponding to resonance frequencies of approximately 15, 25, 35, 45, 55, and 65 MHz). Changes in the normalised resonance frequency (Δ*F* = Δ*f_i_*/*i*) and the dissipation factor (Δ*D*) of the fifth overtone (*i* = 5) are presented throughout. All other overtones provided comparable information.

The thickness of dense protein and lipid layers was estimated from the QCM-D frequency shift using the Sauerbrey equation, as ℎ = −𝐶Δ𝐹/𝜌, with the mass-sensitivity constant *C* = 18.0 ng/(cm^2^ Hz). The film density was assumed to be 𝜌 = 1.1 g/cm^3^ for proteins, and 1.0 g/cm^3^ for lipids, reflecting the solvated nature of the films to a good approximation. We verified that Δ𝐷/−Δ𝐹 ≪ 0.4 × 10^−6^ Hz^-1^, to ascertain films are sufficiently rigid for the Sauerbrey equation to provide reliable film thickness estimates (Reviakine et al., 2011).

### Spectroscopic ellipsometry (SE) for quantification of HC1-PTX3 interaction stoichiometry

SE measurements were performed in situ in a custom-built open cuvette (100 μL volume) with glass windows, on silicon wafers, at room temperature with a spectroscopic rotating compensator ellipsometer (M-2000 V; J.A. Woollam). The ellipsometric angles Δ and Ψ were acquired over a wavelength range from *λ* = 370 to 999 nm, at an angle of incidence of 70°. Samples were directly pipetted into the cuvette, and homogenised by a magnetic stirrer, located at the bottom of the cuvette, for 5 s after sample injection; for the remainder of the sample incubation time, the stirrer was turned off. Excess sample was rinsed away by flowing working buffer through the cuvette; this was assisted by a flow-through tubing system and a peristaltic pump (Ismatec) operated at a flow rate of 5 mL/min; during the rinsing phases, the stirrer was turned on to ensure homogenisation and maximise exchange of the cuvette content.

Data were analysed with the software CompleteEASE (J.A. Woollam) to extract the thickness (*d*_BMF_) and refractive index (*n*_BMF_) of the biomolecular film from the ellipsometric angles, as described in detail elsewhere (Kirichuk et al., 2023). Areal mass densities (*AMD*) and molar surface densities (Γ) of proteins were then calculated using DeFeijter’s equation (Richter et al., 2014), as *AMD* = *d*_BMF_ (*n*_BMF_ - *n*_Sol_) / (d*n*/d*c*) with refractive index increment d*n*/d*c* = 0.18 cm^3^/g and *n*_Sol_ the solution refractive index, and Γ = *AMD* / *M*_W_ with *M*_W_ the protein molecular mass.

### Analytical ultracentrifugation of HC1 and PTX3

Velocity sedimentation AUC analyses was performed on SEC-purified proteins using an XL-A ultracentrifuge (Beckman Coulter, Fullertin, CA, λ = 230 nm) equipped with an An50Ti 8-hole rotor fitted with two-sector epon-filled centrepieces with quartz glass windows. The pH was adjusted to pH 5.5, 6.5 or 7.4 for the respective experiments. Velocity AUC was carried out at 98550 × *g* for HC1-a/s (0.1 mg/mL), and 64800 × *g* for PTX3 (0.1 mg/mL) scanning every 90 s until sedimentation was reached. Sedimentation coefficients were derived from the experimental data using Sedfit (Brautigam, 2015).

## Acknowledgements

We thank Dr Dan Maskell for help with cryo-EM grid preparation and screening (Astbury Biostructure Laboratory, University of Leeds). We also thank Dr Andy Howe (eBIC) for technical support and advice on data collection and Dr Dan Claire for access to Chameleon at eBIC (BI22724-33 and BI29338-1) via Rapid Access Mode. We gratefully acknowledge Dr Matteo Stravalaci and Sonia Valentino for assistance with production of the PTX3 proteins. Crystallography data were collected on beamline I04-1 at Diamond Light Source (proposal MX17773-68). SAXS data were collected on beamline B21 at Diamond Light Source (MX17773-23; MX24447-67). We thank Dr Changjiang You and Prof Jacob Piehler (Osnabrück University, Germany) for providing DODA-(Ni^2+^-NTA)_3_ lipid analogues, and Dr. Steffen Frey and Prof. Dirk Görlich (Max Planck Institute for Multidisciplinary Sciences, Göttingen, Germany) for providing polyhistidine-tagged green fluorescent protein (H_14_-GFP). We also acknowledge Andrea Doni (Humanitas Research Hospital, Italy) and Ruby Law (Monash University, Australia) for helpful discussions early in the project.

## Author contributions

Conceptualisation: A.J.D., A.I., R.P.R.; Data curation: A.J.D, A.S., R.P.R.; Formal analysis: C.B., H.B., T.A.J., C.W.L., M.P.L.C., R.P.R., A.S., M.S., X.Z.; Funding acquisition: C.B., A.J., J.F., A.I., A.M., R.P.R; Investigation: H.B., R.C., L.D., R.J.D., J.F., A.I., T.A.J., C.W.L., M.P.L.C., A.R.E.R., A.S., M.S., X.Z.; Methodology: C.B., H.B., A.J.D., A.I., T.A.J., C.W.L., R.P.R, A.R.E.R., A.S., M.S., X.Z.; Project administration: A.J.D., R.P.R; Resources: A.J.D., J.E., A.I., R.P.R.; Supervision: C.B., A.J.D., R.P.R; Validation: A.J.D., R.P.R, A.S. Visualization: C.W.L., A.J.D., L.D., T.A.J., R.P.R., A.S., M.S., X.Z.; Roles/Writing - original draft: A.J.D., A.I., R.P.R., A.S.; and Writing - review & editing: all authors.

