## Supplementary material for "The structural organisation of pentraxin-3 and its interactions with heavy chains of inter-α-inhibitor regulate crosslinking of the hyaluronan matrix": `S

A.J.D. is a co-founder and employee of Link Biologics Limited, and A.J.D and R.J.D. are both shareholders in the company. Link Biologics is developing a biological drug based on human TSG-6.

Keywords: Cryo-EM; Heavy Chains of Inter-α-inhibitor; Interaction Analysis; Hyaluronan; ITIH1; Pentraxin-3

| **Nomenclature in paper** | **Description** | **Schematic** Protein region shown in purple |
|---|---|---|
| **PTX3** | Full-length, octameric, PTX3. Generated from CHO cells. Used in cryo-EM studies. | 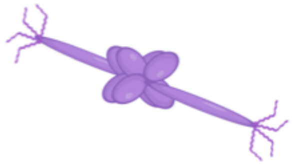 |
| **OCD_PTX3** | Octameric C-terminal domain of PTX3; i.e., comprising 8 C_PTX3 (pentraxin) domains. | 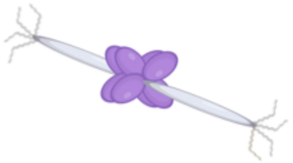 |
| **TND_PTX3** | Tetrameric N-terminal domain; i.e., comprising 4 N_PTX3 domains. | 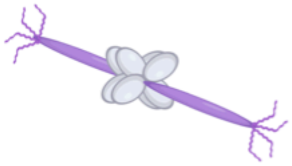 |
| **A48-PTX3** or **D48-PTX3** | Polymorphic variants of PTX3 (SNP: rs3816527). Made in HEK Expi293F (HEK) cells. Used in interaction analyses. See Figure 8 for position of residue 48 on structure. | 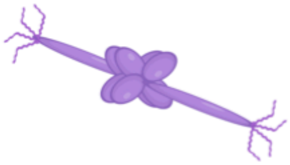 |
| **Half-PTX3** | C317S,C318S mutant made in HEK cells. Replacement of disulphide bonding cysteines in C-terminal region with serines generates PTX3 comprised of 4 protomers. Used in SAXS and interaction analyses. | 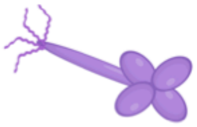 |
| **Half-PTX3-H$_7$** | C317S,C318S mutant with 316-GCCVGGG-322 sequence replaced by 7 His residues. Made in HEK cells. Used in SAXS and interaction analyses. | 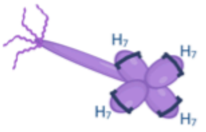 |
| **Δ1-24_PTX3** | Truncated PTX3 where the first 24 amino acids of the N-terminal domain (residues 18-41) are removed. Produced in HEK cells. Used in SAXS and interaction analyses. | 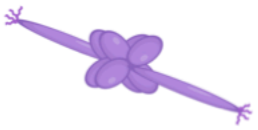 |
| **C_PTX3** | Individual C-terminal (pentraxin) domain of PTX3. Produced in CHO cells. Used in X-ray crystallography. | 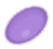 |

**Figure S1. Schematic of the PTX3 constructs and nomenclature used in this work.**

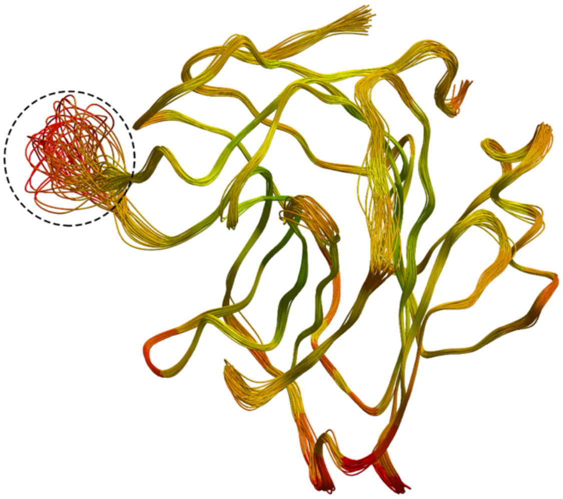

**Figure S2. Ensemble refinement analysis of the PTX3 pentraxin domain.** Ensemble refinement (Burnley *et al*., 2012), a method that uses a combination of refinement and MD simulations, was used to analyse the crystal structure of the C_PTX3 monomer. The 34 models generated are overlayed and colour coded by temperature factor (i.e., from green (least flexible) to red (most flexible)). The region of highest flexibility (dashed circle) corresponds to the loop containing the vicinal cysteines (C317 and C318) that are required for the formation of octamers.

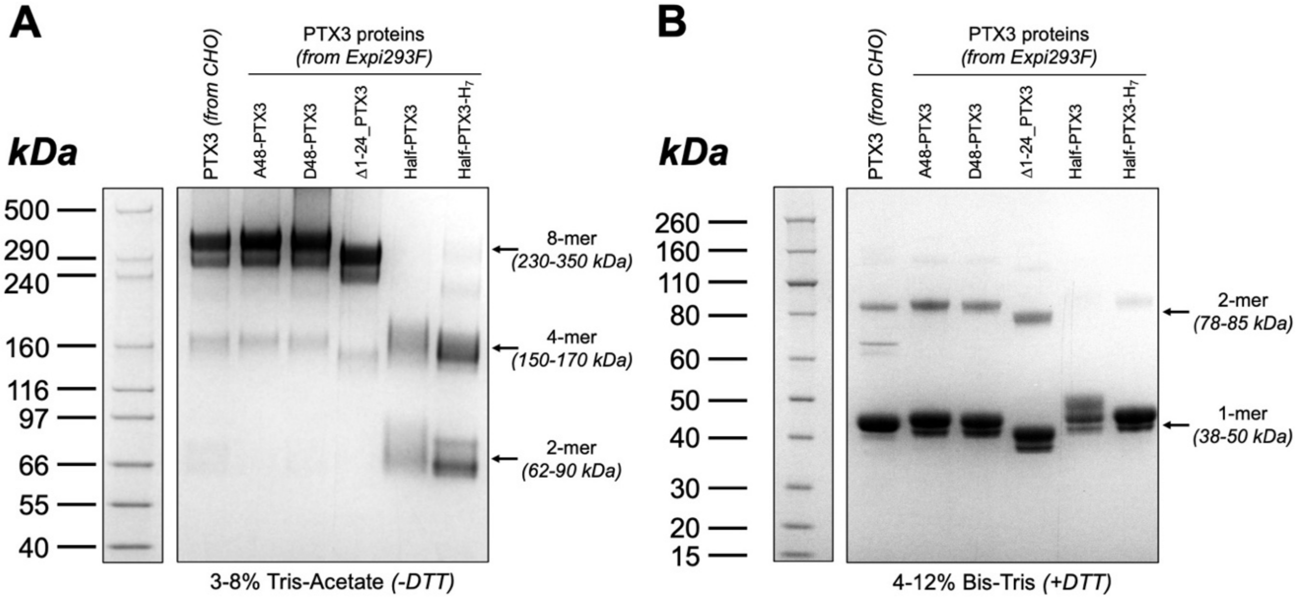

**Figure S3. SDS-PAGE analysis of the PTX3 constructs used in the study.** Full-length human PTX3 from CHO cells and full-length human A48-PTX3 and D48-PTX3 variants, the Δ1-24_PTX3, Half-PTX3 and Half-PTX3-$H_7$ constructs (all made in HEK Expi293F cells) were purified from the conditioned media of transfected cells using a combination of immunoaffinity and size exclusion chromatography. The homogeneity of these preparations was assessed by SDS-PAGE whereby 2 µg/lane of each protein were run on either 3-8% Tris-acetate (**A**) or 4-12% Bis-Tris (**B**) gels, in non-reducing (*-DTT*) and reducing *(+DTT*) conditions, respectively, followed by Coomassie Blue staining. Representative gels of three independent experiments are shown. The position of bands corresponding to monomers (1-mer) and oligomers (2-mer, 4-mer and 8-mer) of PTX3 proteins, and the corresponding ranges of apparent molecular weight, are indicated on the right-hand side of the gels. While the protein preparations are very pure, there is heterogeneity in glycosylation giving rise to the doublets in **A** and **B.** Variance in the disulphide bonds formed by cysteine residues in the N-terminal domain between protomers (Inforzato et al., 2010) leads to a mixture of dimers and tetramers for 'Half-PTX3' constructs in **A**. Incomplete reduction leads to the appearance of dimers in **B**.

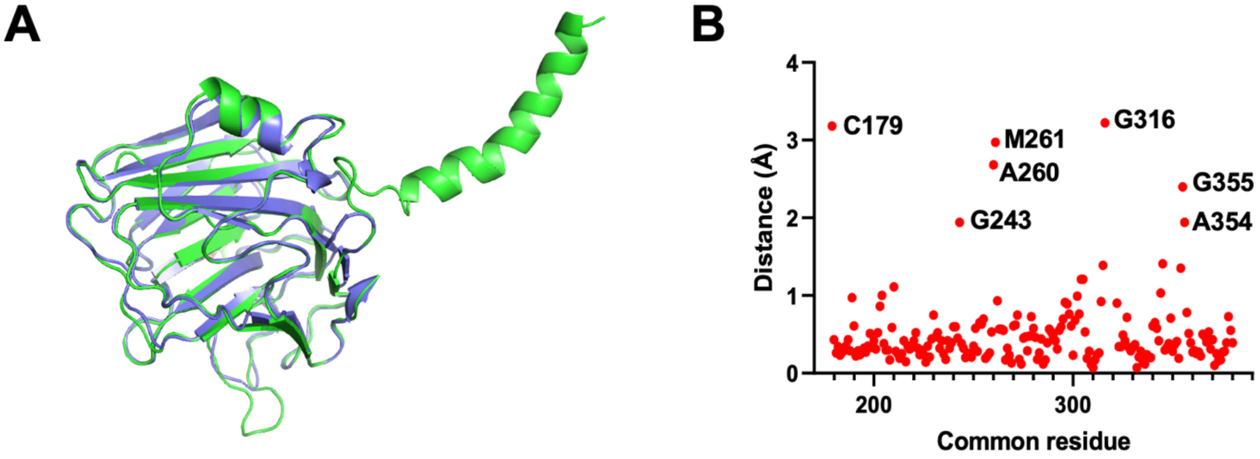

**Figure S4. Structural alignment of the crystal and cryo-EM structures of PTX3.** (**A**) Overlay of the monomeric asymmetric unit of the crystal structure for C_PTX3 (blue) on the cryo-EM model of PTX3 (green) demonstrates that the two structures are highly similar. (**B**) Plot to show the inter-residue distances between common (equivalent) residues of the crystal and cryo-EM structures, showing a high correlation in atomic positions where the backbone RMSD is 0.704 Å. Residues where there is a small divergence in the two structures are indicated on **B**; i.e., due to conformational perturbation of loops, associated with packing of C_PTX3 monomers into an octameric assembly.

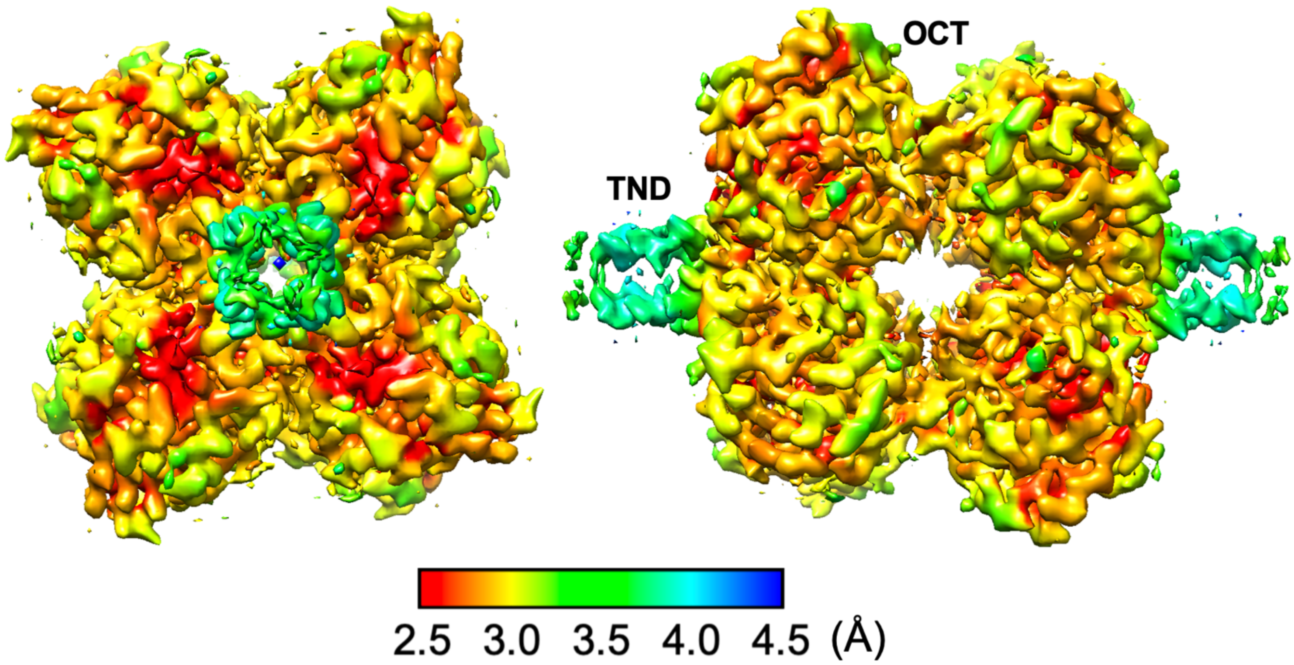

**Figure S5. Local resolution cryo-EM map.** A heat map visualised in CryoSPARC showing the resolution in angstroms (Å) for each voxel of the final cryo-EM map for PTX3. The resolution values at each atom position were generated in Chimera. The right-hand map is rotated by 90˚ around the *y*-axis relative to the left-hand map. The Octameric C-terminal Domain (OCD) of PTX3 has a resolution, that is mostly ≤3.0 Å, whereas the Tetrameric N-terminal Domain (TND) is of lower resolution, being mostly resolved to 3.5-4.0 Å.

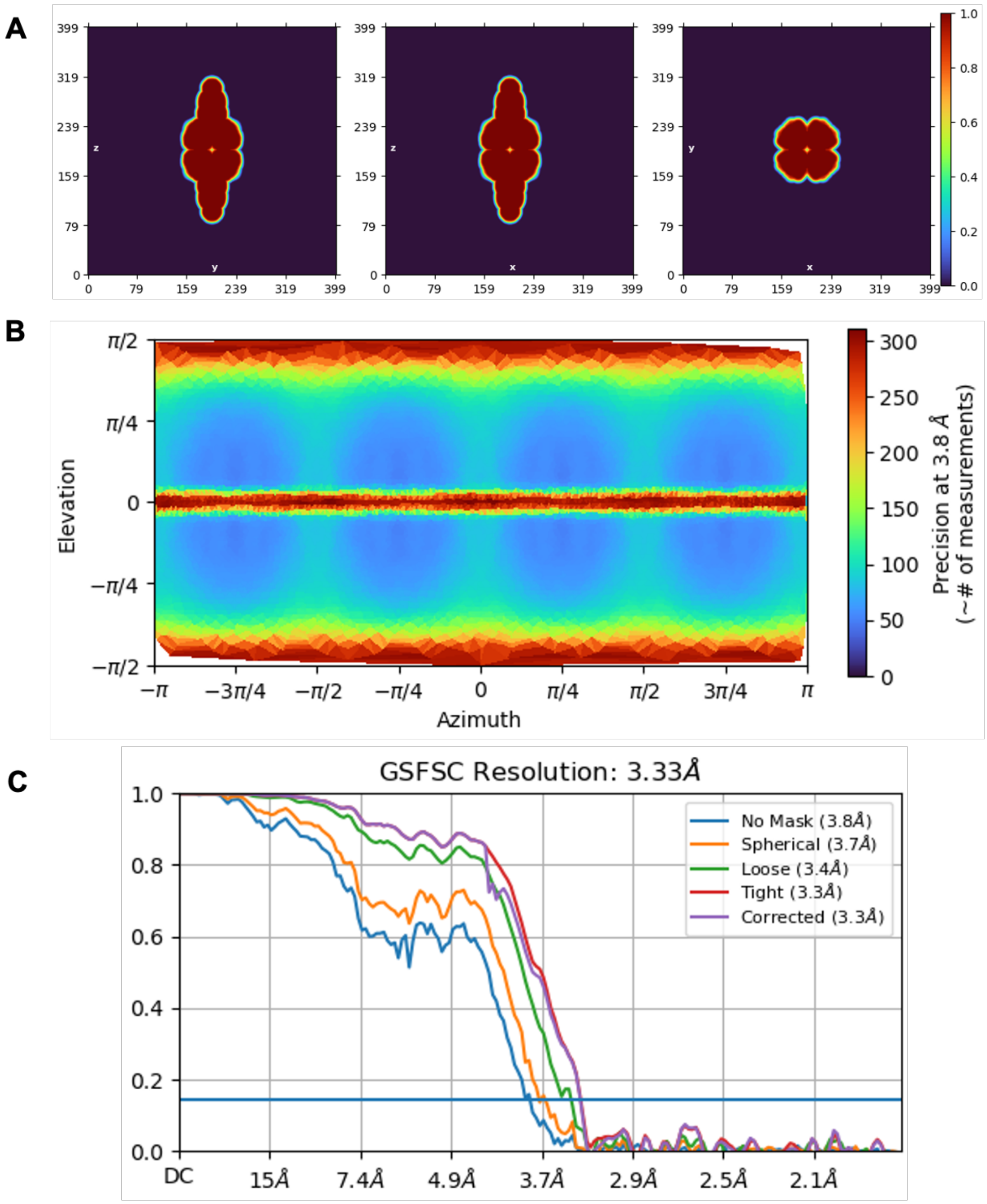

**Figure S6. Validation of cryo-EM map.** CryoSPARC was used to validate the cryo-EM map generated in this study. (**A**) Input mask used for local resolution jobs. (**B**) Plot to show the angular distribution of particles. (**C**) The Fourier Shell Correlation (FSC) showing a global resolution estimation (GSFSC) of 3.33Å.

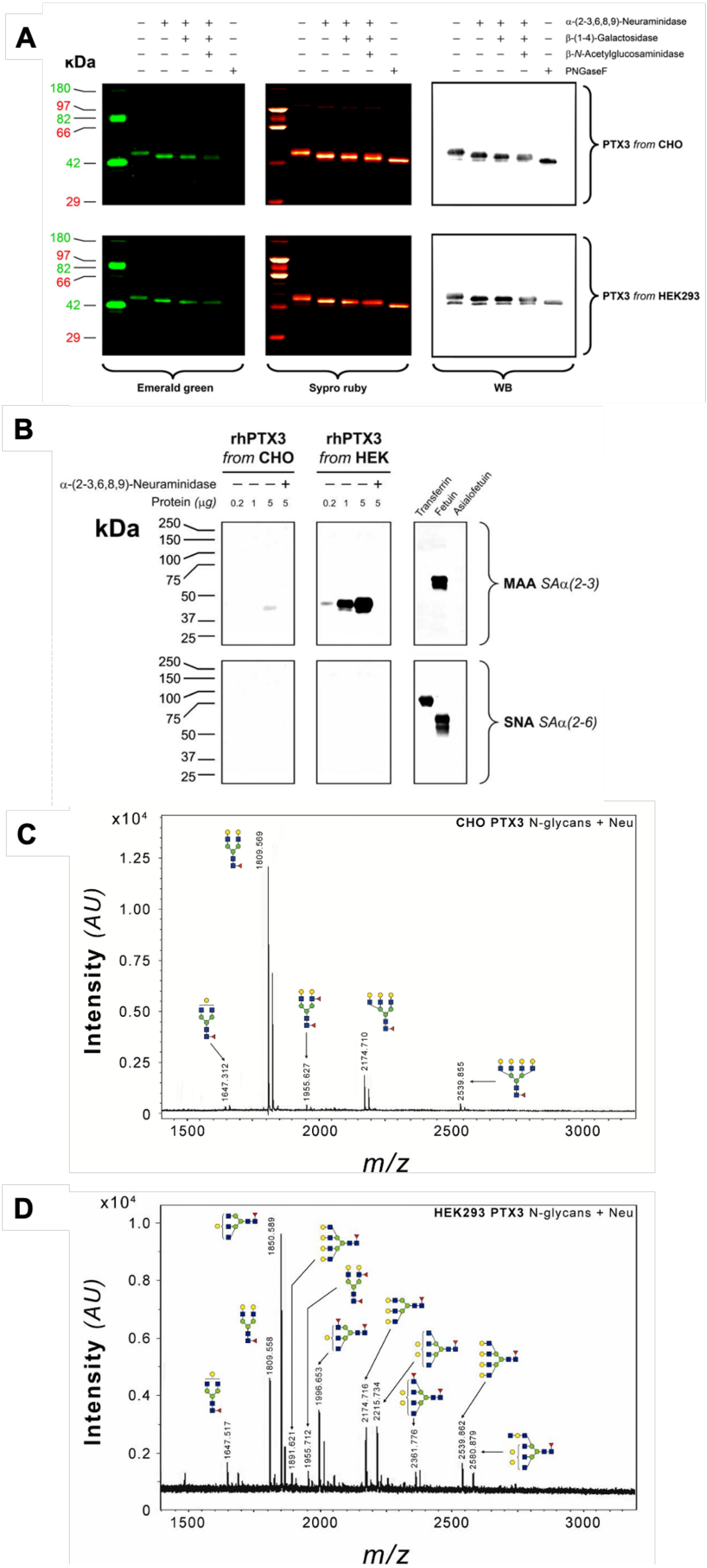
A
κDa
α-(2-3,6,8,9)-Neuraminidase
β-(1-4)-Galactosidase
β-N-Acetylglucosaminidase
PNGaseF
180
97
82
66
42
29
PTX3 from CHO
PTX3 from HEK293
Emerald green
Sypro ruby
WB
B
rhPTX3 from CHO
rhPTX3 from HEK
α-(2-3,6,8,9)-Neuraminidase
Protein (μg)
0.2 1 5 5
Transferrin
Fetuin
Asialofetuin
kDa
250
150
100
75
50
37
25
MAA SAα(2-3)
SNA SAα(2-6)
C
CHO PTX3 N-glycans + Neu
x10^4
1.25
1.00
0.75
0.50
0.25
0
1809.569
1647.312
1955.627
2174.710
2539.855
Intensity (AU)
1500
2000
2500
3000
m/z
D
HEK293 PTX3 N-glycans + Neu
x10^4
1.0
0.8
0.6
0.4
0.2
0
1850.589
1809.558
1647.517
1891.621
1955.712
1996.653
2174.716
2215.734
2361.776
2539.862
2580.879
Intensity (AU)
1500
2000
2500
3000
m/z

**Figure S7. Characterisation of the glycans from recombinant PTX3 proteins expressed from CHO and HEK Expi293F cells.** (**A**) CHO- and HEK Expi293F-derived PTX3 proteins (top and bottom panel, respectively) were incubated with linkage specific exoglycosidases or PNGaseF (that catalyses deamidation of N-glycosylated Asn residues with the removal of the N-glycans) and separated by SDS-PAGE on 10% Tris-Glycine gels under reducing conditions (2 µg/lane). This was followed by glycoprotein (Emerald green, i.e., Pro-Q Emerald 300) or general protein (Sypro ruby) fluorescent staining. In separate experiments, 20 ng/lane aliquots of the same materials were transferred onto PVDF membranes (after SDS-PAGE) and probed by Western blotting (WB) with a rabbit anti-human PTX3 polyclonal antibody. Protein deglycosylation can be seen as both an increase of electrophoretic mobility and decrease of Emerald green fluorescence. (**B**) Extent and linkage specificity of terminal sialylation were assessed by lectin blotting. The PTX3 preparations from CHO and HEK Expi293F cell were incubated in the presence or absence of $\alpha$-(2-3,6,8,9)-neuraminidase (that catalyses hydrolysis of $\alpha$-(2-3,6,8,9)-linked sialic acid; SA), separated on 10% Tris-Glycine SDS-PAGE gels (0.2, 1 and 5 µg/lane) and transferred onto nitrocellulose membranes. The membranes were probed with either *Maackia amurensis* (MAA) or *Sambucus nigra* (SNA) lectins that recognize $\alpha$-(2-3)- and $\alpha$-(2,6)-SA, respectively. Transferrin, fetuin and asialofetuin were used as positive and negative controls, respectively. (**C,D**) Oligosaccharides were released from the polypeptide backbone of CHO- (**C**) and HEK Expi293F- (**D**) derived PTX3 by PNGase F digestion, incubated with $\alpha$-(2-3,6,8,9)-neuraminidase (Neu; to remove negatively charged SA), purified with non-porous graphitised carbon microcolumns and analysed by MALDI-MS on a Bruker Reflex III MS system operated in reflectron positive ion mode by adaptation of a previous protocol (Morelle and Michalski, 2007). This allowed recording of sodiated pseudomolecular ions that were assigned to glycoform structures using the GlycoMod tool (https://web.expasy.org/glycomod/). Symbols used for the monosaccharide components of the recorded oligosaccharides are indicated. Gels and MS spectra shown in **A** to **D** are representative of at least three independent experiments with the same results.

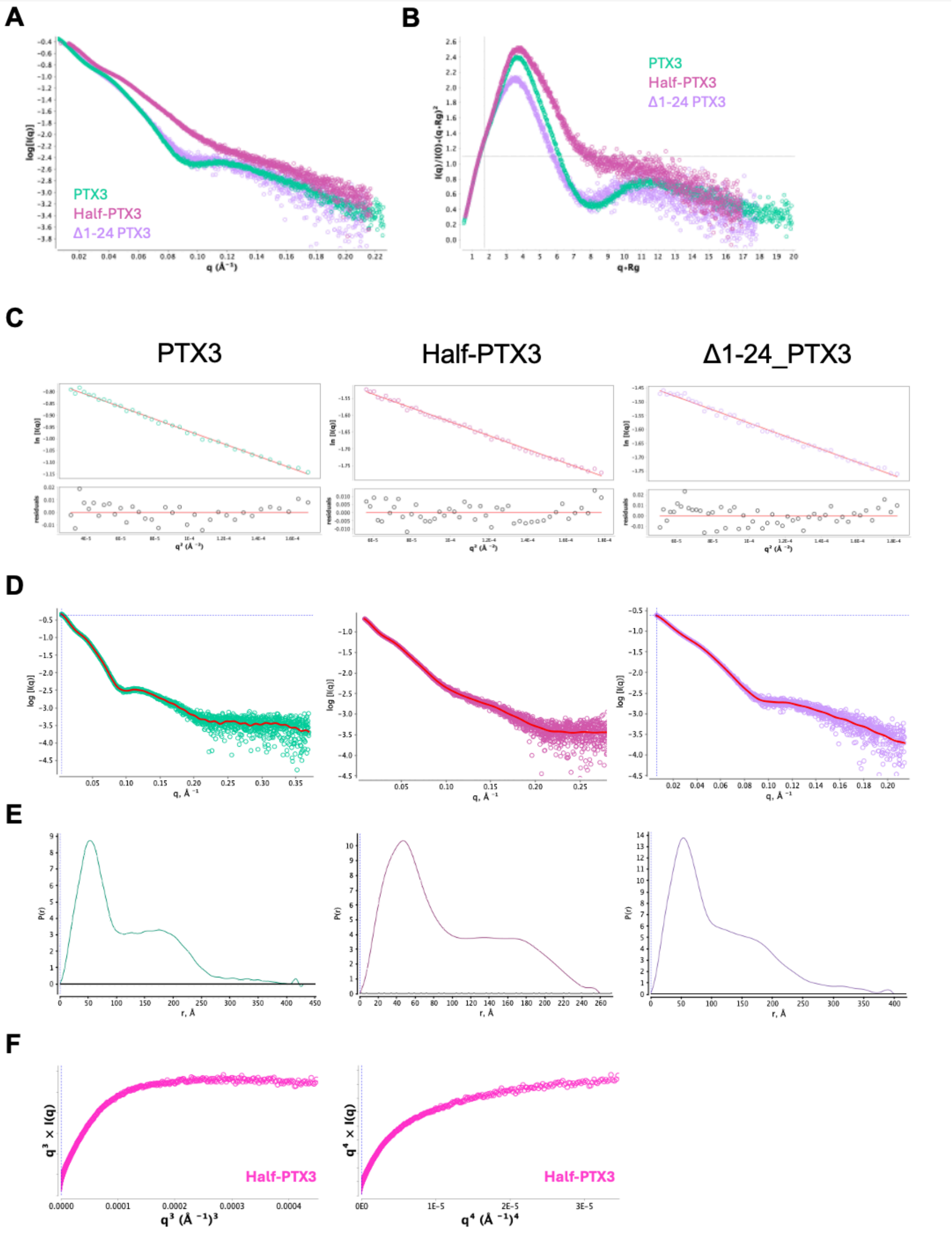

**Figure S8. SAXS analysis of PTX3 proteins**. SAXS data for PTX3 (green), Half-PTX3 (pink) and Δ1-24_PTX3 (purple) were analysed using ScÅtterIV: (**A**) Overlay of the $\log_{10}$ plots; (**B**) Overlay of the dimensionless Kratky plots; (**C**) Analysis of the Guinier region of the data; (**D**) Fits of the data to obtain the $d_{max}$ (see Table S1) and (**E**) *P*(*r*) distance distributions; (**F**) SIBYLS ($q^3$) and Porod-Debye ($q^4$) plots for Half-PTX3.

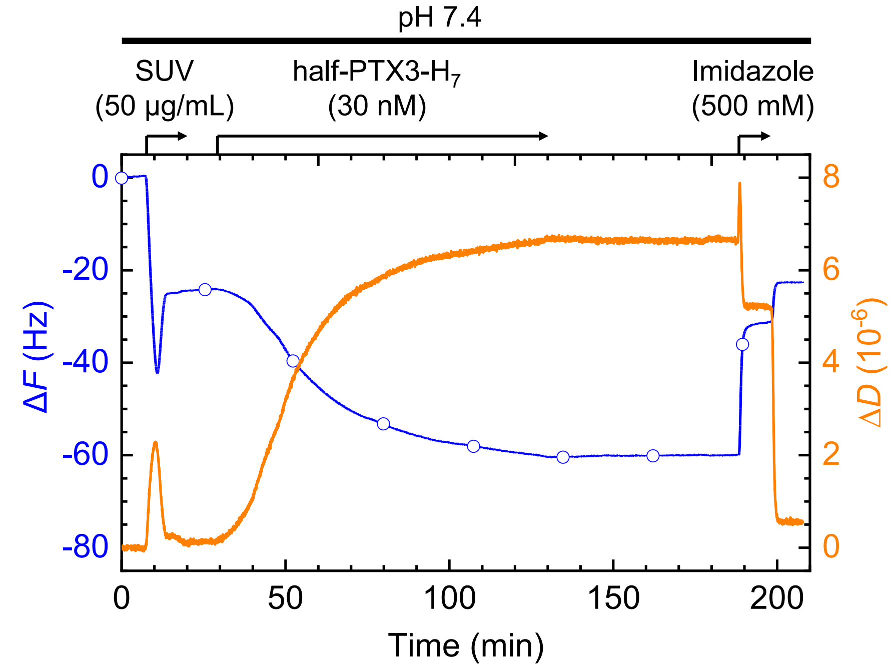

**Figure S9. Validation of appropriate surface anchorage of half-PTX3-$H_7$ to supported lipid bilayers using QCM-D.** QCM-D resolves changes in the mass and mechanical properties of biomolecular films at the surface-liquid interface. To a first approximation, a negative frequency shift ($\Delta F$) correlates with a mass increase, which comprises the surface bound biomolecules as well as hydrodynamically coupled solvent, and a positive dissipation shift ($\Delta D$) correlates with the increased softness of the biomolecular film. Thanks to the combination of both parameters, QCM-D provides information about the morphology of biomolecular films and their interactions with analytes. $\Delta F$ (blue line with open circles) and $\Delta D$ (orange line) data are from the $5^{th}$ overtone ($i$ = 5). The initial part (0 to 25 min) is a representative example of SLB formation from SUVs containing a mix of 99.5 mol-% DOPC and 0.5 mol-% DODA-($Ni^{2+}$-NTA)$_3$. The two-phase response with local extrema in $\Delta F$ and $\Delta D$ is characteristic for vesicles initially adhering intact, and then gradually rupturing and coalescing into a confluent SLB (Richter et al., 2006). The final frequency shift ($\Delta F$ = -25 ± 1 Hz) indicates a film of 4.5 ± 0.2 nm thickness, as expected for SLBs. The low final dissipation shift ($\Delta D < 0.5 \times 10^{-6}$) is characteristic of SLBs of good quality, with minimal surface coverage of un-ruptured vesicles. The latter part (25 to 210 min) demonstrates site-specific anchorage of half-PTX3-$H_7$ in HBS (pH 7.4), via binding of its His-tags to $Ni^{2+}$-NTA, which is stable to rinsing in buffer, but can be rapidly and fully eluted with 500 mM imidazole. Arrows above the traces represent the start and duration of sample incubations, with concentrations as indicated; during remaining times, HBS alone was flowed over the sensor surface. Changes in $\Delta F$ and $\Delta D$ upon exchange of imidazole from/to HBS do not reflect any alterations on the surface but result from a change in the viscosity and density of the surrounding solution owing to the presence/absence of imidazole.

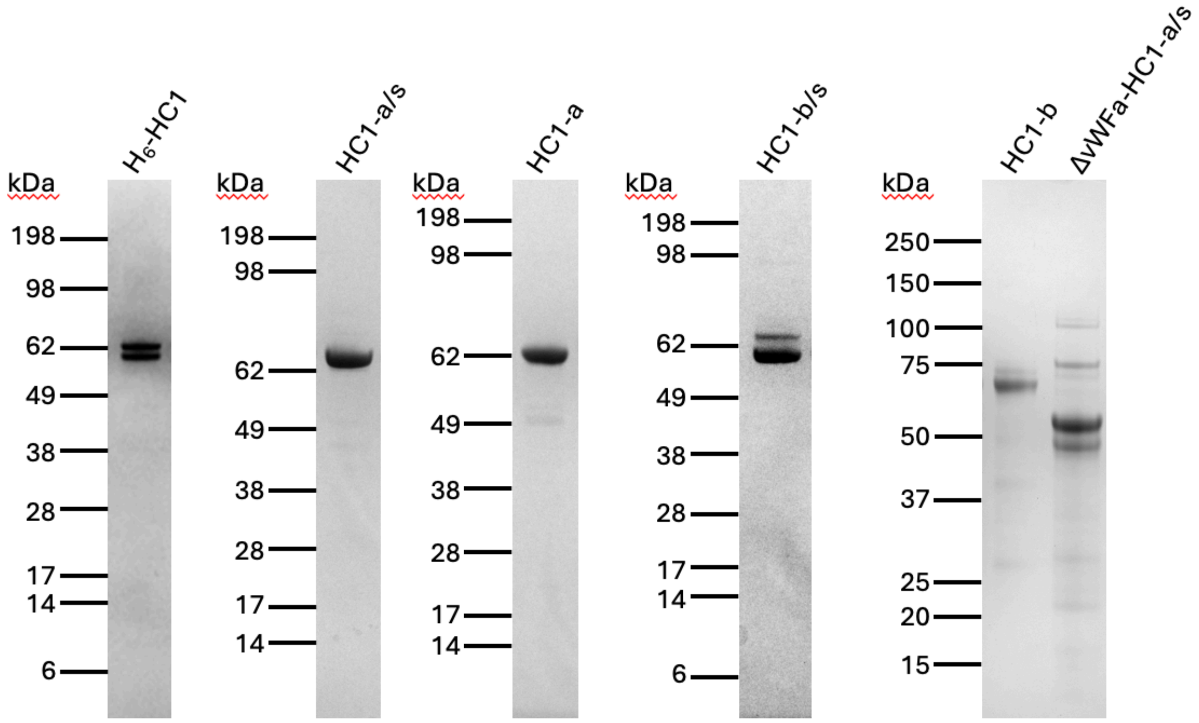

**Figure S10. SDS-PAGE analysis of the HC1 constructs used in the study.** SDS-PAGE under reducing conditions for $H_6$-HC1 (1 µg), HC1-a/s (1.5 µg), HC1-a (1.5 µg), HC1-b/s (1 µg), HC1-b (2 µg) and ∆vWFa-HC1-a/s (2 µg). The first four proteins were run on separate gels shortly after being purified; MW (to left of gel lane) are based on See Blue Plus2 MW size markers (Invitrogen). The last two proteins were run on the same gel (with Precision Plus Protein Standards (Bio-Rad)) more than 1 year after being made, explaining the degradation in ∆vWFa-HC1-a/s; mass spectrometry at time of production (Table 2) showed that the protein was within 2.5 Da of theoretical mass. The individual images of the gels have been adjusted to give a similar level of background colour, but no other image manipulation was performed.

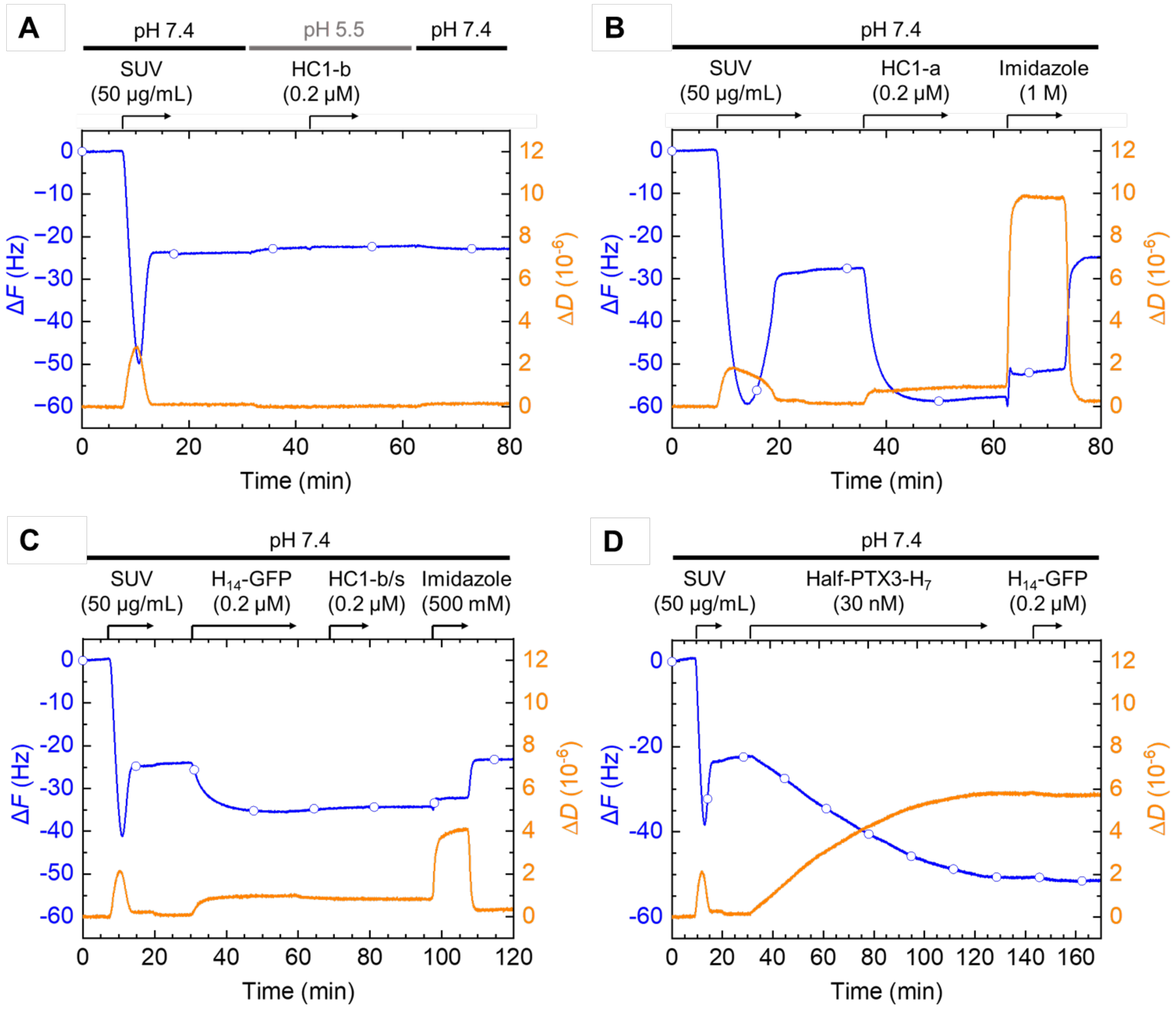

**Figure S11. Validation of specific binding of HC1 constructs to SLB-anchored Half-PTX3-$H_7$ by QCM-D**. Δ*F* (blue line with open circles) and Δ*D* (orange line); data are from *i* = 5 ($5^{th}$ overtone). (**A**) HC1-b does not bind to plain DOPC SLBs at pH 5.5. (**B**) HC1-a binds to SLBs with DODA-($Ni^{2+}$-NTA)$_3$; such binding can be fully eluted with imidazole, indicating it involves $Ni^{2+}$ chelation, most likely via some of the 28 histidine residues in this construct. Such non-specific binding was mitigated against by avoiding any excess of DODA-($Ni^{2+}$-NTA)$_3$ lipids in the SLBs beyond what was required for Half-PTX3-$H_7$ anchorage. (**C**) HC1-b/s binding to SLBs containing 0.5 mol-% DODA-($Ni^{2+}$-NTA)$_3$ can be eliminated by blocking with a His-tagged protein (here green fluorescent protein with an N-terminal $H_{14}$ tag ($H_{14}$-GFP (Ananth et al., 2018). This demonstrates that the strategy to mitigate non-specific HC1 binding works well. (**D**) Half-PTX3-$H_7$ effectively blocks all $Ni^{2+}$-NTA binding sites in SLBs containing 0.5 mol-% DODA-($Ni^{2+}$-NTA)$_3$; following saturation of PTX3 binding, $H_{14}$-GFP was incubated to test for any residual binding capacity of the SLB. Virtually no binding was observed. Conditions used: in **A**, HBS (10 mM HEPES, pH 7.4, 150 mM NaCl) and MBS (10 mM MES, pH 5.5, 150 mM NaCl); in **B**-**D**, HBS. Arrows above the graphs represent the start and duration of sample incubations (with concentrations as indicated); during remaining times buffer alone was flowed over the sensor surface. SUVs contained 0 mol-% (**A**), 5 mol-% (**B**), or 0.5 mol-% (**C**, **D**) DODA-($Ni^{2+}$-NTA)$_3$ in a DOPC background. Note that changes in Δ*F* and Δ*D* upon switching from HBS to imidazole (in **B**, **C**) are caused by the increase in viscosity and/or density of the solution upon addition of high imidazole concentration, and do not reflect any changes of the surface.

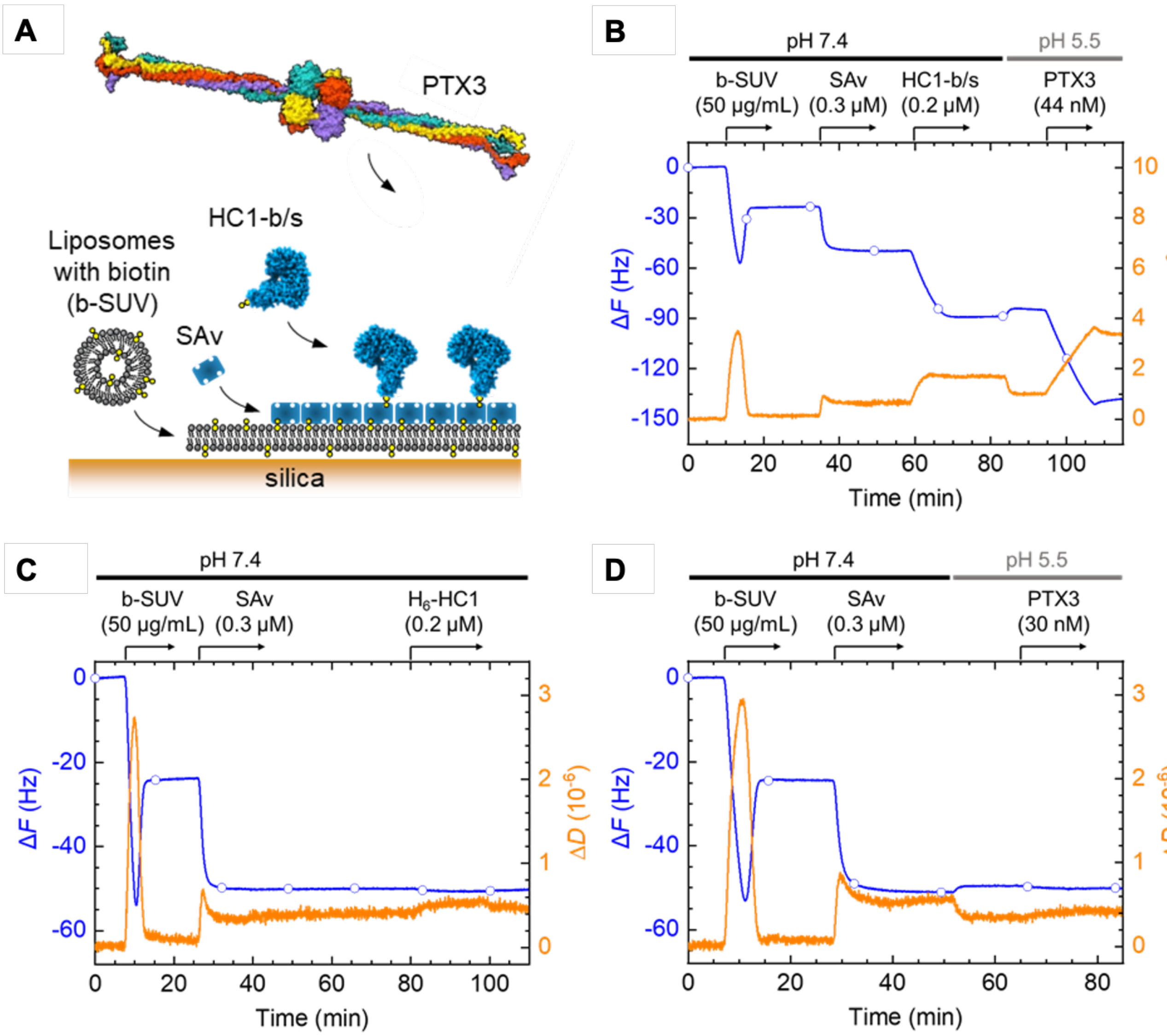

**Figure S12. Evidence of HC1-PTX3 interactions in an 'inverted' binding assay.** (**A**) Schematic of the inverted binding assay: HC1-b/s, anchored via a C-terminal biotin/Strep II (b/s) tag to a dense monolayer of streptavidin covering a biotin-presenting supported lipid bilayer (SLB) (made from SUVs with 95 mol-% DOPC and 5 mol-% DOPE-cap-biotin), interacts with PTX3 analyte in the solution phase. (**B**) Representative QCM-D data (Δ*F*: blue line with open circles; Δ*D*: orange line; from 5th overtone (*i* = 5)) from the binding assay set up in **A**. Responses for SLB and streptavidin monolayer formation are as previously established (Migliorini *et al.*, 2014). HC1-b/s binding (starting at 60 min) saturates at -Δ*F* = 40 Hz, indicating formation of a ~7 nm thick monolayer, a value consistent with the size of HC1 (~11 nm; (Briggs et al., 2020)). HC1-b/s binding is stable upon rinsing in HBS (at 73 min). A shift in pH from 7.4 to 5.5 (at 83 min) induces a small yet clear decrease in Δ*D* and an increase in Δ*F*. This indicates HC1 film compaction at acidic pH, consistent with the AUC analysis (Figure 5C, Figure S13). PTX3 binding (starting at 93 min) engenders pronounced QCM-D responses, as expected for a large protein. PTX3 binding is essentially stable upon rinsing in pH 5.5 (at 106 min). (**C**) QCM-D data demonstrating site-specific anchorage via the b/s tag in this assay: HC1 lacking a b/s tag (though carrying a N-terminal $His_6$ tag) does not bind to SAv. (**D**) QCM-D data demonstrating specificity of PTX3 binding to HC1 at pH 5.5: there is no appreciable binding of PTX3 to bare SAv under this condition. Buffer conditions used: MBS (10 mM MES, pH 5.5, 150 mM NaCl) or HBS (10 mM HEPES, pH 7.4, 150 mM NaCl). Arrows above the graphs represent the start and duration of sample incubations (with concentrations as indicated); during remaining times buffer alone was flowed over the sensor surface; horizontal lines at the top indicate the solution pH (pH 7.4 – black, pH 5.5 – grey).

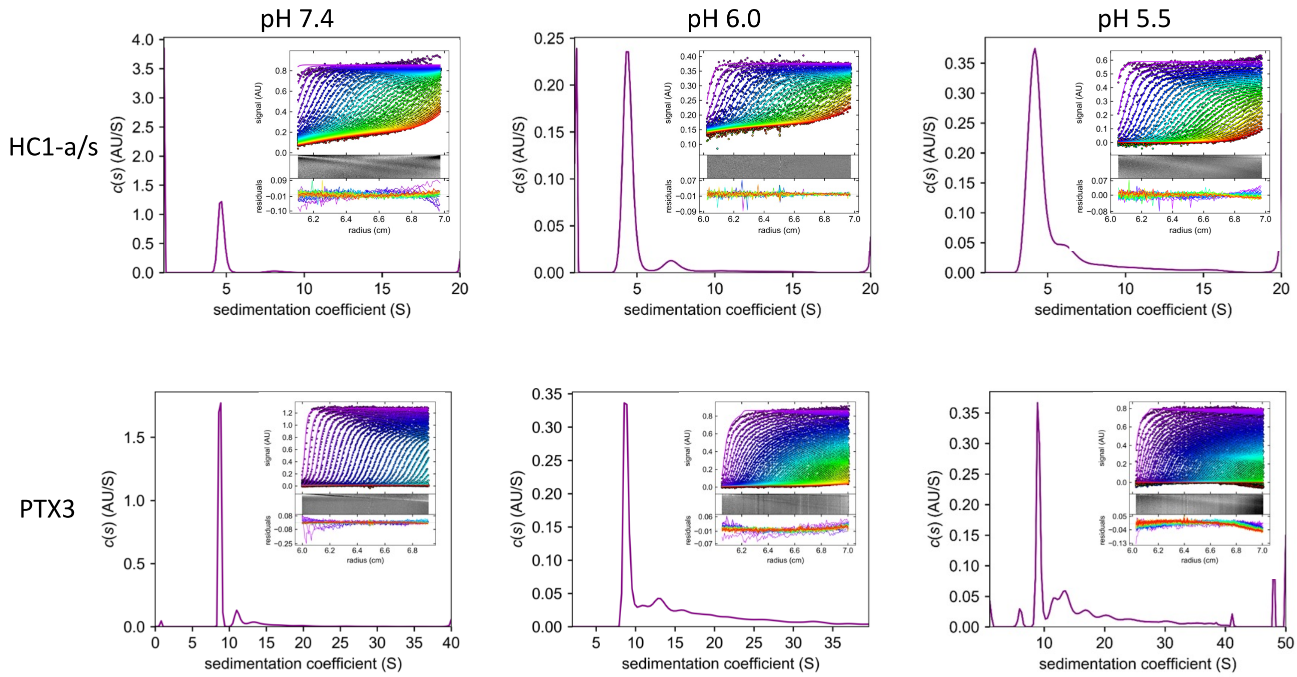

**Figure S13. Sedimentation velocity analysis of HC1 and PTX3.** Main images show the relative distribution of sedimentation coefficient *c*(s) for HC1-a/s (top panels) and PTX3 (bottom panels) at pH 7.4 (left-hand panels), pH 6.0 (middle panels) and pH 5.5 (right-hand panels). The data were fit using a single ideal *c*(s) model (Brown and Schuck, 2006). The insets show the raw data (every 3$^{rd}$ scan) with the model fit and the residuals as both a bitmap image and a graph of the deviation.

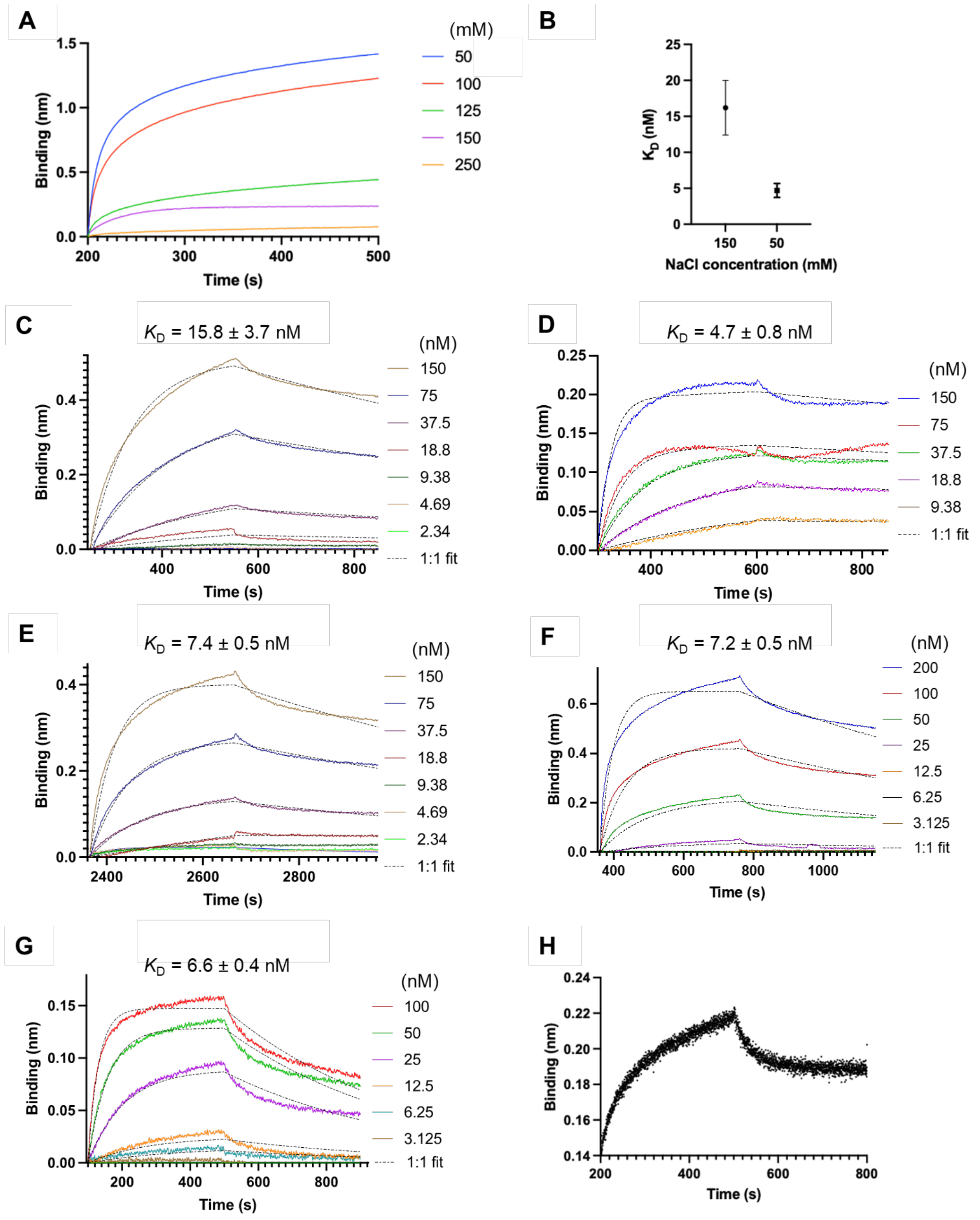

**Figure S14. Salt-strength dependence of the HC1-PTX3 interaction at pH 5.5.** (**A**) BLI data for the binding of A48-PTX3 (100 nM) analyte to HC1-a/s ligand, at a range of NaCl concentrations (colour coded as indicated). **(B)** Binding affinities determined by BLI analysis for 150 mM and 50 mM NaCl concentrations (mean ± SD for *n* = 3 technical replicates on the same HC1-a/s surface). (**C**) BLI data for the binding of A48-PTX3 analyte (at a range of concentrations, colour coded as indicated) to HC1-a/s ligand in MBS (10 mM MES, pH 5.5, 150 mM NaCl) as described in Figure 5D. (**D**) BLI data for the binding of A48-PTX3 analyte (at a range of concentrations, colour coded as indicated) to HC1-a/s ligand in a low salt buffer (10 mM MES, pH 5.5, 50 mM NaCl). (**E**) BLI data for the binding of D48-PTX3 analyte (at a range of concentrations, colour coded as indicated) to HC1-a/s ligand in MBS. (**F**) BLI data for the binding of Δ1-24_PTX3 analyte (at a range of concentrations, colour coded as indicated) to HC1-a/s ligand in MBS. (**G**) BLI data for the binding of PTX3 analyte to ΔvWFA_HC1-a/s ligand in MBS. (**H**) BLI data for the binding of A48-PTX3 analyte to ΔvWFA_HC1-a/s ligand, in the presence of 2.5 mM EDTA in MBS; while only a single concentration was analysed, the presence of a binding event demonstrates that the interaction is metal ion independent.

**Table S1. SAXS analysis of PTX3 proteins.**

| | **PTX3** | **Half-PTX3** | **Δ1-24_PTX3** |
|---|---|---|---|
| Guinier Analysis | | | |
| $I(0)$ ($cm^{-1}$) | 0.492 | 0.242 | 0.261 |
| $R_g$ (Å) | 88.19 | 77.90 | 83.72 |
| Low $q$ ($Å^{-1}$) | 0.0059 | 0.013 | 0.007 |
| High $q$ ($Å^{-1}$) | 0.225 | 0.220 | 0.215 |
| $P(r)$ analysis | | | |
| $I(0)$ ($cm^{-1}$) | 0.500 | 0.242 | 0.281 |
| $R_g$ (Å) | 97.33 | 79.38 | 95.34 |
| $d_{max}$ (Å) | 430 | 260 | 400 |
| Analysis from Primus | | | |
| $R$(g) (Å) | 86.49 | 70.52 | 79.31 |
| $d_{max}$ (Å) | 497 | 323 | 429 |
| $q_{min}$ | 0.08093 | 0.09929 | 0.08826 |
| $q_{max}$ | 0.30012 | 0.25003 | 0.20004 |
| $M_W$ $V_c$ (kDa) | 323.873 | 181.359 | 331.621 |
| $M_W$ MoW (kDa) | 380.271 | 221.504 | 384.929 |
| $M_W$ Bayesian Inference (kDa) | 318.450 | 208.000 | 318.450 |

**Table S2. Cryo-EM acquisition parameters for Datasets 1 and 2.**

| | **Dataset 1** (EMD-19717) | **Dataset 2** (for 3DVA analysis) |
|---|---|---|
| **Microscope** | Titan Krios (eBIC; BI22724-33) | Titan Krios 2 (University of Leeds) |
| **Detector** | Falcon 4 with Selectris energy filter | Falcon 4 (counting mode) |
| **Accelerating Voltage (kV)** | 300 | 300 |
| **Pixel size (Å)** | 0.72 | 0.86 |
| **Nominal Mag** | 16,500× | 96,000× |
| **Dose ($e^-$/pix/sec)** | 5.00 | 7.14 |
| **Exposure (s)** | 4.4 | 5.0 |
| **Total dose (e/Å$^2$)** | 40.00 | 48.24 |
| **Defocus range (μm)** | -1.0 to -2.2 | -1.0 to -2.2 |
| **Spherical aberration (mm)** | 2.7 | 2.7 |
| **Images collected** | 12,114 | 11,700 |

**Table S3. BLI replicate data**.

| | **Ligand immobilised** | **Analyte in solution** | $K_D$ **(nM)** | $k_a$ **(1/Ms)** | $k_{dis}$ **(1/s)** | $\chi^2$ |
|---|---|---|---|---|---|---|
| Repeat 1 | HC1-a/s | PTX3 | 5.17 | 2.10E+05 | 1.08E-03 | 0.6804 |
| Repeat 2 | HC1-a/s | PTX3 | 4.67 | 2.11E+05 | 9.85E-04 | 0.7238 |
| Repeat 3 | HC1-a/s | PTX3 | 4.71 | 1.74E+05 | 8.19E-04 | 2.5762 |
| **Average ± SD** | | | **4.85 ± 0.23** | **1.98E+05** | **9.62E-04** | |
| Repeat 1 | HC1-a/s | Δ1-24_PTX3 | 6.68 | 1.28E+05 | 8.52E-04 | 7.9808 |
| Repeat 2 | HC1-a/s | Δ1-24_PTX3 | 7.19 | 1.37E+05 | 9.86E-04 | 4.4701 |
| Repeat 3 | HC1-a/s | Δ1-24_PTX3 | 7.63 | 1.35E+05 | 1.03E-03 | 0.9921 |
| Average **± SD** | | | **7.17 ± 0.39** | **1.33E+05** | **9.56E-04** | |
| Repeat 1 | ΔvWFA_HC1-a/s | PTX3 | 6.11E-09 | 3.08E+05 | 1.88E-03 | 0.6359 |
| Repeat 2 | ΔvWFA_HC1-a/s | PTX3 | 6.76E-09 | 3.40E+05 | 2.30E-03 | 0.5673 |
| Repeat 3 | ΔvWFA_HC1-a/s | PTX3 | 6.87E-09 | 3.82E+05 | 2.63E-03 | 0.582 |
| **Average ± SD** | | | **6.58 ± 0.34** | **3.43E+05** | **2.27E-03** | |
| Repeat 1 | HC1-a/s | A48-PTX3 | 10.6 | 7.12E+04 | 7.57E-04 | 1.0274 |
| Repeat 2 | HC1-a/s | A48-PTX3 | 17.9 | 7.44E+04 | 1.33E-03 | 1.0884 |
| Repeat 3 | HC1-a/s | A48-PTX3 | 18.9 | 8.29E+04 | 1.57E-03 | 0.6781 |
| **Average ± SD** | | | **15.8 ± 3.7** | **7.62E+04** | **1.22E-03** | |
| Repeat 1 | HC1-a/s | D48-PTX3 | 6.83 | 1.36E+05 | 9.26E-04 | 1.1523 |
| Repeat 2 | HC1-a/s | D48-PTX3 | 7.55 | 1.45E+05 | 1.09E-03 | 1.0808 |
| Repeat 3 | HC1-a/s | D48-PTX3 | 7.86 | 1.43E+05 | 1.12E-03 | 0.8754 |
| **Average ± SD** | | | **7.41 ± 0.44** | **1.41E+05** | **1.05E-03** | |
| Repeat 1 | HC1-a/s | PTX3* | 4.66 | 9.54E+04 | 4.45E-04 | 0.0959 |
| Repeat 2 | HC1-a/s | PTX3* | 3.76 | 1.80E+05 | 6.75E-04 | 0.9838 |
| Repeat 3 | HC1-a/s | PTX3* | 5.70 | 1.36E+05 | 7.76E-04 | 0.4634 |
| **Average ± SD** | | | **4.7 ± 0.79** | **1.37E+05** | **6.32E-04** | |

*= in 50 mM NaCl-containing buffer as opposed to 150 mM used for all other experiments.

**Table S4. Cryo-EM structure refinement statistics for 8S50.pdb and EMD-19717.**

| | |
|---|---|
| **Reconstruction** | |
| Number of particles | 23,457 |
| Symmetry | D4 |
| Map resolution (Å; FSC threshold = 0.143) | 3.33 |
| Resolution range (Å; min, 25th percentile, median, 75th percentile, max) | 1.978, 2.901, 3.688, 5.628, 10.553 |
| Map sharpening B-factor (Å$^2$) | -137.2 |
| **Refinement** | |
| Model resolution (Å; FSC threshold = 0.5) | 3.41 |
| FSC model (Å; 0, 0.143, 0.5) | 3.15, 3.28, 3.41 |
| Overall Biso (Å$^2$) | 20 |
| CC (mask, box, volume, peaks, main chain, side chain | 0.8, 0.78, 0.8, 0.75, 0.81, 0.79 |
| **Model quality** | |
| Clashscore | 10 |
| Ramachandran (%; favoured, allowed, outliers) | 91.21, 8.79, 0.00 |
| Sidechain Outliers (%) | 0.1 |
| C-beta outliers (%) | 0 |
| Molprobity score | 2.02 |
